# An organotypic neocortical slice culture for studying neuroglial interactions

**DOI:** 10.64898/2026.05.15.725074

**Authors:** Kieran P. Higgins, Viktor A.B. Al Naqib, Pierre Mayo, Bastiaan Lodder, Takahiro Masuda, Lukas Amann, Marco Prinz, Maarten H.P. Kole

## Abstract

Organotypic slice cultures (OSCs) are widely used to study cellular properties in a functional and developmental tissue context. With the recent advent of transgenic mouse lines and viral tools we postulated that OSCs may enable the study of multicellular glial and neuroglial interactions in development, as well homeostatic and pathological conditions. Here, we made mouse cortical OSCs and used markers for oligodendroglial, microglial states and neuronal types between 1 to 28 days in vitro (DIV). The OSC was characterized by in-vivo like cortical layering, including layer 5 pyramidal neurons and produced highly robust synchronized period bursts resembling Up- and Down states. Glial cells showed a strong cortical layer- and time-dependent development pattern: in the first week (DIV 1–7), slicing-related debris clearance and developmentally restricted sparse oligodendroglial myelination created an environment with highly phagocytic, non-homeostatic microglia (assessed with CD68 and purinergic receptor P2Y12, respectively). Between DIV 14 and 21, however, slices showed stereotypical cortical myelin patterns and the emergence of a homeostatic microglia phenotype while exhibiting continued phagocytosis. Furthermore, live two-photon imaging and morphometric analyses revealed highly ramified microglia and myelinated axons with compact myelination, exceeding lamellae count compared to age-matched in vivo axons. Lastly, from DIV 28 and onwards, myelin integrity became impaired and associated with phagocytic microglia. Together, the results indicate that between DIV14 and 21 cortical OSCs are well suited for live imaging of homeostatic and activity-dependent neuron-glia interactions, bridging the gap between in vivo investigations and primary cell cultures.

## Introduction

The evolutionary appearance of the central nervous system (CNS) is inextricably linked to the advent of glia, providing multiple and diverse contributions to neuronal functions, both in health and disease ^1^. Oligodendrocytes produce multilamellar myelin sheaths around neuronal axons to enable the rapid and temporally precise propagation of action potentials and provide a critical source for activity-dependent metabolic support ^2,3^. Microglia, the brain’s tissue macrophages and one of the first lines of self-defense against pathogens, are critically implicated in fine-tuning neural circuitry during development ^4–8^. In addition, microglia also engage with other glial cells and, for example, play an important role in maintenance of oligodendroglial myelination ^9,10^. Such bidirectional interactions between glial cells act to facilitate proper neuronal function, and dysfunctions of neuron-glia or glia-glia interactions are emerging as key processes in driving neurodegenerative disease ^11,12^ or the pathogenesis of neuropsychiatric disorders ^13–16^ .

An understanding of the molecular and cellular properties of neuron-glia interactions requires the integrative study of these interactions in the native complex biological context of brain tissue. While *in vivo* studies are the ‘gold standard’ in brain research, they come with the drawback of restricted ability for manipulations and limited spatiotemporal resolution. Light scattering in tissue limits imaging in deeper layers of the cortex and the ability to resolve the fine glial and neuronal processes, at which neuron-glia interactions are often taking place – near the diffraction limit of light microscopy. To bridge the gap between intact brain tissue and dissociated cell cultures, and image at the resolution of neuron-glia contact sites, organotypic brain slice cultures offer a preparation that preserves neuronal connectivity and cytoarchitecture and offers an *in-vivo* like environment ^17–19^. Importantly, slice cultures harbor all brain-resident cell types, with microglial RNA expression profiles closely resembling those of the intact brain, and better suitability to study (patho)physiological roles of microglia when compared to traditional 2D cultures ^20,21^. Furthermore, due to their culturing at an interface state, both local application of agents such as viruses, as well as systemic manipulations through the culturing medium can be performed ^22^. This high degree of manipulability, combined with ease of access to lower cortical layers for imaging and electrophysiological recordings potentially could make cortical organotypic cultures a valuable tool for investigating neuro-microglia and neuron-oligodendrocyte interactions. While slice cultures have been well characterized for the hippocampus ^22–24^, cerebellum ^25,26^ and the neocortex ^25,27–29^ there is a paucity in the knowledge on the development and presence of glia cells.

Here, we examined the layer-specific development of microglia, oligodendrocytes and neurons in organotypic slices cultures (OSCs) from the mouse neocortex. We utilized the microglia-specific *Hexb*^tdTomato/tdTomato^ (here referred to as *Hexb*^tdTom^) reporter line ^30,31^ and in combination with viral tools labelled oligodendrocytes and their myelin membranes ^32^. We report the ability to follow detailed morphological properties and spatiotemporal patterns of glia development across cortical layers for 4 weeks in culture. OSCs from the *Rbp4*-Cre line showed a preservation of cortical layering in deeper layers. Using confocal, 2-photon, and electron microscopy, as well as electrophysiological recordings, we uncovered key developmental timepoints defined by microglial state changes and increasing number of mature myelinating oligodendrocytes. Between 14 and 21 days in vitro (DIV) the neocortical OSC contains homeostatic microglia and robust axonal myelination, with both similarities and differences compared to in vivo myelination features. Together, our data reveal a robust platform to study the molecular properties of neuron-glia and glia-glia interactions in neocortical circuits.

## Results

### Slice protocol and viability

To investigate the development of neuronal and glial cell interactions, cortical organotypic slice cultures (OSCs) were prepared from 4–5 day old mouse pups from either C57BL/6*, Rbp4-*Cre or *Hexb*^tdTom^ strains (**Figure 1A**, see Methods for detail). In part based on previous methods ^33,34^, mice were sacrificed by decapitation with scissors, and the brain was extracted and placed in chilled dissection solution ice cold dissection buffer. Brains were cut down the midline and then sectioned via McIlwain Tissue Chopper to obtain 300 µm thick coronal slices. Slices were quickly transferred to membrane inserts and kept at 35 °C and 5% CO_2_ for the duration of the culturing period. By nature, the preparation of organotypic slice cultures requires the traumatic act of slicing the brain, which may result in damage to brain areas intended for investigation ^19^. To quantify the viability of the cortex, we used propidium iodine (PI) to label dead and degenerating cells in culture and following fixation counterstained with the DNA marker 4′,6-diamidino-2-phenylindole (DAPI) and visualized the cell death from DIV 1 to 28 (**Figure 1B** and **C**). Immunofluorescence imaging and subsequent image analysis revealed a significantly larger fraction of compromised cells in subcortical structures compared to the cortex one day after slicing (*P* = 0.0002, **Figure 1D**). Although cellular degeneration substantially and significantly reduced for all subsequent time points after DIV 1 (*P* > 0.353, **Figure 1D**), we noticed that further into the culturing period, at DIV 21 and 28, regions of the cortex exhibited bands of PI-positive cells (**Figure 1B**). Resolving cellular degeneration by cortical depth, we observed a substantial amount of dying cells in the center of the cortex, reflecting putative layers (L) 4 and 5, at DIV 21 (**Figure 1E**). This broad distribution of degeneration narrowed and moved upwards along the cortical column a week later, with superior layers showing marked cell death at DIV 28. These observations indicate that, following the slicing-induced cleanup of cellular debris, a steady state of cell health encompasses the entire cortex, which then leads to decreased cell health in localized patches of weeks 3 to 4.

**Figure 1.**
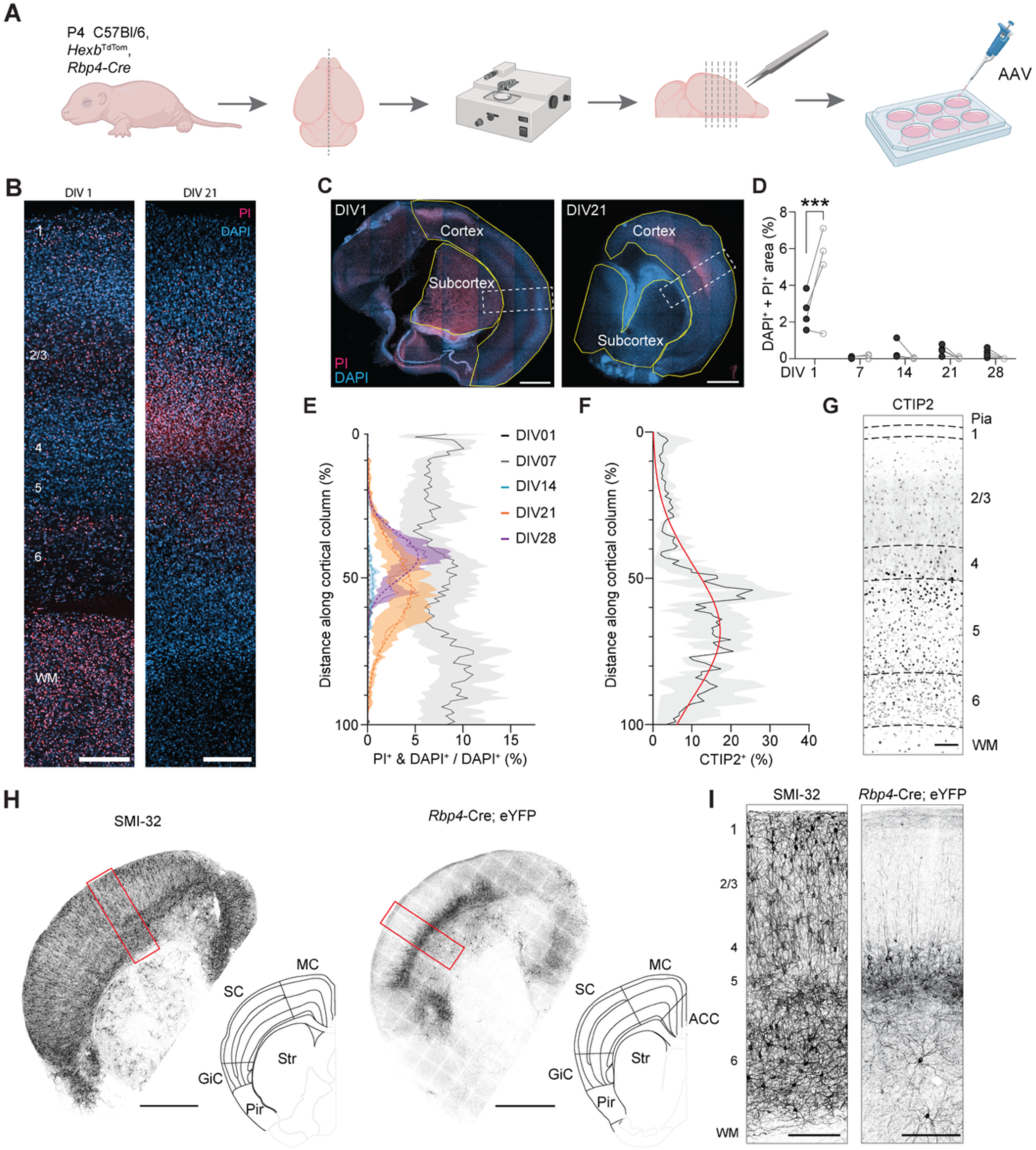
Neocortical organotypic slice cultures preserve cortex and its stereotypical architecture over weeks in culture. **A.** Schematic of the OSC preparation strategy. **B.** Single *z*-plane apotome images of cortical columns from DIV 1 (left) and DIV 21 (right) OSCs with DAPI (cyan) and PI (magenta). Scale bars, 200 µm. **C.** Representative apotome images of OSCs at DIV 1 (left) and DIV 21 (right). Outlines of quantified areas (cortex/subcortex) in yellow, insets in (B) as dashed boxes. Scale bars, 1 mm. **D.** Time course of cellular degeneration in cortex (black) and subcortex (grey) shows substantial cell death happening predominantly in extracortical areas at DIV 1; (two-way ANOVA, DIV \*\*\*\**P* < 0.0001 , region *P* = 0.2096, interaction \*\**P* = 0.0014; Tukey’s multiple comparisons \*\*\**P* < 0.001, n = 4 slices per DIV). **E.** Laminar distribution and time course of degenerating cells in OSCs along the cortical column from pia (0%) to white matter (100%). Means from n = 4 slices indicated by solid lines, shaded area is the SEM; Gaussian fits for DIV 7, 14, 21, 28 (R^2^ = 0.14, 0.34, 0.36, 0.54, respectively) as dashed lines. **F.** CTIP2^+^ area running from pia (0%) to the white matter (100%), Gaussian fit shown in red (R^2^ = 0.44); *n* = 3 slices. **G.** Representative image of a DIV 14 OSC stained for CTIP2, with lines indicating the cortical lamina (WM, white matter). Scale bar, 100 µm **H.** Left, SMI-32 staining at DIV 21; right, eYFP staining at DIV21 with *Rbp4*-Cre-dependent expression of ChR2-eYFP; red box indicating inset area. Insets show matched sections from the Allen Reference Atlas ^35^ – Mouse Brain (P56) to the right of each slice. Motor cortex (MC), somatosensory cortex (SC), gustatory-insular cortex (GiC), anterior cingulate cortex (ACC) and striatum and piriform area (Str, Pir). Scale bar, 1 mm **I.** Left, crop of H, showing the cortical lamina distribution of SMI-32. Right, crop of H, depicting band of *Rbp4*-Cre-driven expression in L5. Scale bar, 250 µm.

Since cortical viability appeared to vary based on cortical depth, we next sought to investigate the preservation of cortical layering in OSCs. To this end, we conducted immunostainings against CTIP2, a transcription factor found both early in development and in the mature neocortex, specifically expressed in subcortically projecting neurons in the deeper layers ^36,37^. Consistent with the preservation of layering we found that at DIV 14, CTIP2 immunofluorescence staining showed a peak intensity halfway from pia to white matter, in a band at the start of L5, and a gradual decrease in immunoreactivity upon approaching the border of L6 and white matter (**Figure 1F** and **G**). To further study the neurons in OSCs at DIV 14-21, we visualized SMI-32 using immunostainings, which recognizes an epitope of nonphosphorylated neurofilament protein, a marker of a subpopulation of pyramidal neurons localized in L2/3 and L5 of various neocortical regions of the rodent ^38,39^. Interestingly, while deeper layers showed strong SMI-32 staining, the neurofilament protein was also observed in neuronal types across layers from pia to subcortical area in the OSC **(Figure 1H**). Finally, to genetically test the presence of L5 pyramidal neurons, we prepared OSCs from the *Rbp4-*Cre driver line, which labels extra- and intra-telencephalic projecting L5 pyramidal neuron populations ^40^. In line with *in vivo* studies, expression was observed mostly in neurons with cell bodies localized to the deeper layers (**Figure 1I**). Together, these findings indicate that the slicing protocol and slice incubation ensures the survival of the cortex for 2–3 weeks depending on cortical depth, and preserves key cell types in L5, showing broadly conserved stereotypical organization.

### Microglia attain a homeostatic profile during the culturing period

The main phagocytes in OSCs – the microglia – exhibit genetic signatures of inflammatory phenotype in the first week(s) of culturing ^21^, and we next asked if microglia were impacted over time in culture. Using immunohistochemical staining of the cortical column from DIV 1 to 28, we quantified the state in which microglia reside along their phagocytic and homeostatic axes both over time and in the tissue context. In the first week, microglia (demarcated by the marker Iba1) lacked the homeostatic marker P2Y12 while the lysosomal and endosomal marker CD68 showed moderately high expression in ∼20% of the Iba1^+^ area (**Figure 2A-D**). While microglia at DIV 1 appeared amoeboid, they took up a characteristically ramified morphology and were practically devoid of P2Y12 at DIV 7 (**Figure 2B**). In contrast, P2Y12 expression significantly and dramatically increased at DIV 14 to reduce slightly and significantly with time in culture (**Figure 2C**). Concomitantly, CD68 expression at these time points remained statistically unchanged with respect to the first week in culture (**Figure 2D**). During DIV14–21 the microglia resembled a classic homeostatic phenotype, with long, bifurcated processes characterized by P2Y12. In the last week of culture, we observed a trend towards both increased CD68 as well as a decline in P2Y12 expression, hinting at gradual slice degeneration (**Figure 2C** and **D**). Cells from this time point further showed higher expression of CD68 in their somata and processes (**Figure 2B**).

**Figure 2.**
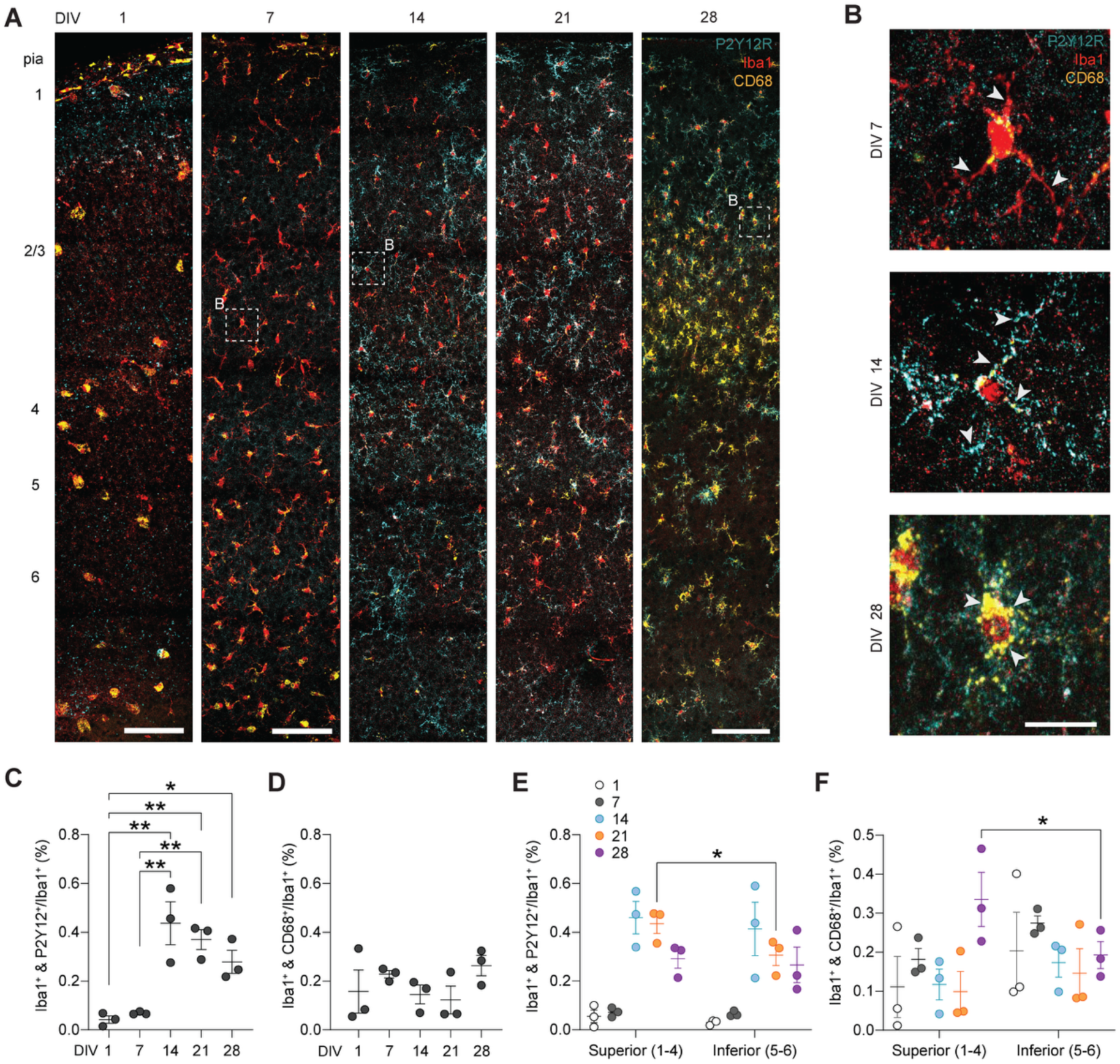
Microglia transition from a phagocytic to a homeostatic profile. **A.** *Z*-projected confocal images depicting developmental timeline of immunostainings for microglial phagocytosis and homeostatic states throughout culture. DIV 1-28, with P2Y12 (cyan), Iba1 (red), and CD68 (yellow). Dashed box insets shown in (B). Scale bar, 100 µm **B.** Representative microglial cells from DIV 7, 14, and 28; crop from (A). Arrows indicate hallmark features at the respective time points; scale bar 20 µm. **C.** Microglial P2Y12 expression emerges at 2 weeks in culture. Iba1^+^ P2Y12^+^ area normalized to Iba1^+^ area, assessed over DIV1–28 (one-way ANOVA *** *P* = 0.0005 with Tukey’s post-hoc test, \*\**P* < 0.0097, \**P* = 0.0404 *n* = 3 slices) **D.** Microglial CD68 remains unchanged throughout culturing. Iba1^+^ and CD68^+^ area normalized to Iba1^+^ area, assessed over DIV 1-28 (one-way ANOVA *P* = 0.3584) **E.** Microglia remain homeostatic for longer in superior layers (1–4) compared to inferior layers (5–6) (two-way ANOVA, DIV \*\*\**P* = 0.0005, layer \**P* = 0.0228, interaction *P* = 0.2566; Šídák’s post-hoc test, \**P* = 0.0367) **F.** CD68 expression of microglia in superior layers significantly increases compared to inferior layers at 4 weeks in culture. (two-way ANOVA, DIV *P* = 0.1190, layer *P* = 0.4715, interaction \**P* = 0.0129; Šídák’s post-hoc test, \**P* = 0.0297)

We noticed that the expression pattern in microglia was nonuniformly distributed across the cortical layers, with for example bands of highly CD68-positive cells in upper layers at DIV 28 (**Figure 2A**). Since cellular degeneration in the OSC showed increased apoptosis in the upper layers (**Figure 1**), we next asked if microglial states were lamina dependent. When averaging fluorescence signals within the superior (1 to 4) and inferior layers (5 to 6) we observed that at DIV21 microglia within the inferior layers expressed P2Y12 less abundantly, in line with a decrease in viability in the deeper layers (*P* = 0.0367, **Figure 2E**). Furthermore, consistent with the time-dependent shift in viability, DIV 28 microglia in the superior layers were significantly more phagocytic when compared to cells in inferior layers (*P* = 0.0297, **Figure 2F**). Additionally, extracortical and white matter-associated microglia were found to have a very high phagocytic load, barely ramified morphologies, and low P2Y12 expression. The findings thus indicate that the cortex is the most viable and homeostatic region of the cultured neonatal brain slice (**Supplemental Figure 1**).

Taken together, these data show that microglia undergo developmental maturation during culturing and their states are layer dependent. Furthermore, the microglial participation in phagocytosis throughout the culturing period does not preclude their ability to express homeostatic markers.

### Morphologies of parenchymal OSC microglia resemble those in vivo

The microglial expression profile in DIV 14–21 suggests a homeostatic state (**Figure 2**). To obtain additional evidence we made live 2-photon images of the morphology using *Hexb*^tdTom^ mice with *z*-stacks encompassing the entire depth of the slice, from the air-exposed top traversing the central parenchyma to the membrane-attached bottom (**Figure 3A**). While we normally constrained our analyses to the parenchyma, we compared cells with the slice border and made quantitative comparisons of microglia morphology by semi-automatically tracing single microglia in L2/3 from of DIV 14–21 OSCs and multiple live fields of view were assembled into a tile scan covering the entire column (**Figure 3B,C**). When comparing microglia from top to the bottom of the slice, differences in ramification were immediately apparent as those cells in the parenchyma were markedly more ramified than at the borders (**Figure 3A** and **B**, **Supplemental Figure 2**). Sholl analysis of cells from the three locations underpinned this observation, where parenchymal microglia showed a markedly higher ramification in comparison to cells from the bottom of the slice (**Figure 3D**). Further, by plotting morphometric parameters obtained from 3D reconstructions we found that parenchymal microglia were significantly more complex than those at the bottom (**Figure 3F-H**). This pertained to the absolute length of arborizations, the number of terminal end points, as well as the number of branch nodes. Microglia at the air-exposed top differed from parenchymal cells in some, but not all parameters, indicating an intermediate state.

**Figure 3.**
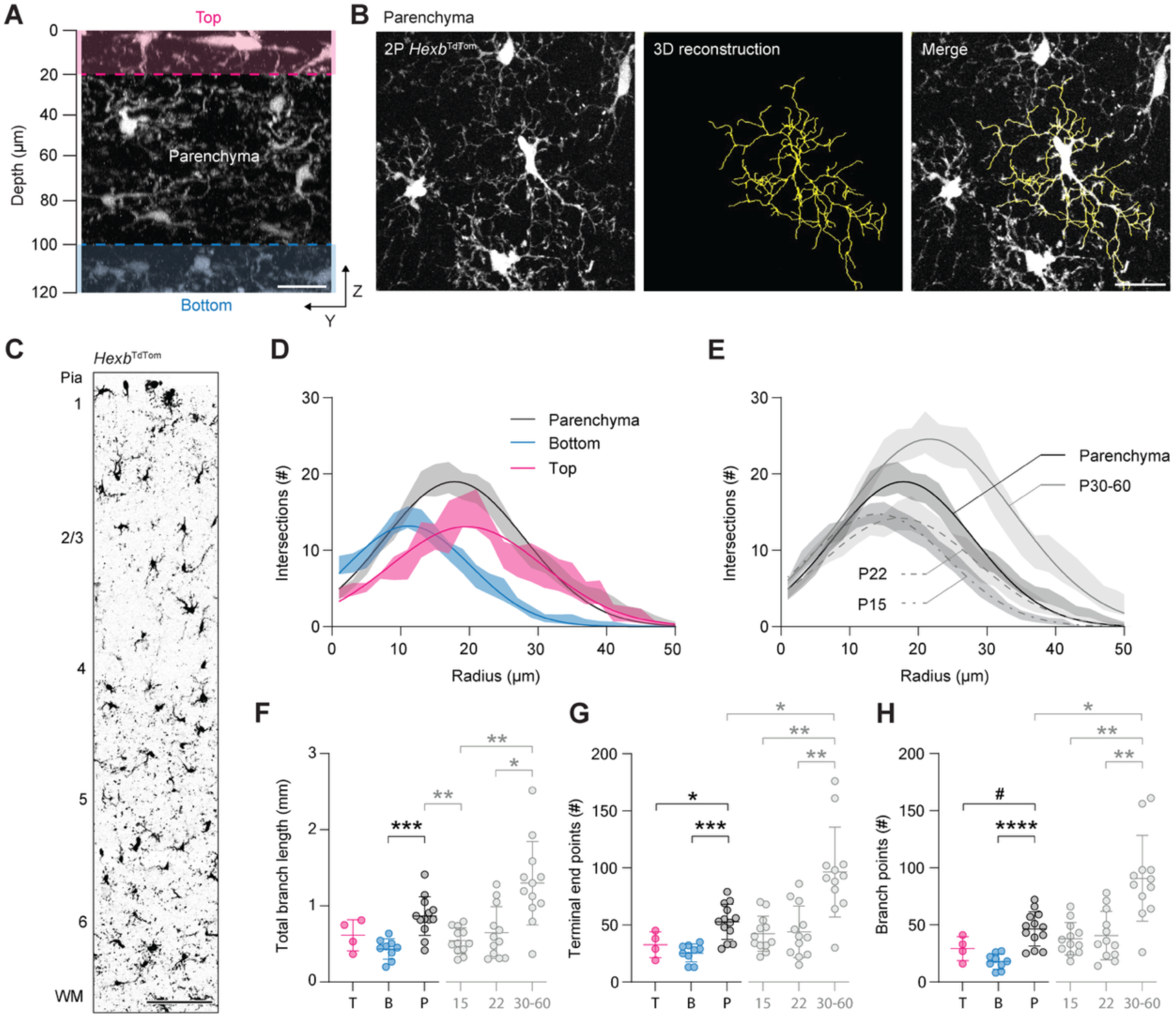
Parenchymal microglia show a homeostatic morphology. **A.** 2-photon Y-Z-projection of along the whole thickness of a DIV 18 *Hexb*^tdT^ OSC. Dotted lines indicate borders between the fringes of the slice (top/bottom) and the parenchyma. Scale bar 20 µm. **B.** Example images of microglia from a *Hexb*^tdT^ OSC at DIV 22 with fluorescence signal (left, grey) morphological reconstruction (center, yellow), and merge (right). Scale bar 20 µm. **C.** 2-photon tile scan of a *Hexb*^tdT^ OSC at DIV 18 showing laminar distribution of microglia, from the pia to the white matter (WM). Scale bar 100 µm. **D.** Sholl analysis of microglia from several locations in OSCs indicates that parenchymal microglia are highly ramified cells. Lines indicating Gaussian fit (R^2^ = 0.7550, 0.7613, 0.7821 for P, T, and B respectively) with SEM of underlying data in shaded area. **E.** Ramification of parenchymal microglia is intermediate between P15/22 and P30-60 *in vivo* microglia. Lines indicating Gaussian fit (R^2^ = 0.7550, 0.7768, 0.6482, and 0.6852 for P, P15, P22, and P30-60, respectively) with SEM of underlying data in shaded area. **F-H.** Microglial morphometric parameters from cells in the parenchyma (P), top (T), and bottom (B) of OSCs, as well as from P(15), P(22), and P(30-60) primary somatosensory cortex *in vivo*. Comparisons between parenchymal and top/bottom microglia on the left and in black, comparisons between parenchymal cells and *in vivo* on the right and in grey (*n*(P) = 13, *n*(T) = 4, *n*(B) = 9, *n*(15) = 12, *n*(22) = 12, *n*(30-60) = 12). **F.** Parenchymal microglia have more extensive total branch lengths than cells at the bottom of OSCs, while they are comparable to microglia from both P22 and P30-60 (left, Welch ANOVA, \*\*\**P* = 0.0002 with Dunnett’s T3 post-hoc test, \*\*\**P* = 0.0002; right, Welch ANOVA ***P = 0.0003 with Dunnett’s T3 post-hoc test, \**P* = 0.0155, \*\**P* < 0.007). **G.** Number of terminal end points for microglia in the parenchyma is comparable to cells from P15/22 *in vivo* but differs significantly from P30-60 *in vivo* microglia or cells from OSC borders (left, one-way ANOVA \*\*\**P* = 0.0002 with Tukey’s post-hoc test, \*\*\**P* = 0.0001, \**P* = 0.0292; right, Welch ANOVA \*\*\*\**P* < 0.0001 with Dunnett’s T3 post-hoc test, \**P* = 0.0170, \*\**P* < 0.0051). **H.** Parenchymal microglia have more bifurcated processes than cells at slice borders and are similar to microglia from P15 and P22 (left, one-way ANOVA \*\*\*\**P* < 0.0001 with Tukey’s post-hoc test, ^#^*P* = 0.0534, \*\*\*\**P* < 0.0001; right, Welch ANOVA \*\*\*\**P* < 0.0001 with Dunnett’s T3 post-hoc test, \**P* = 0.0106, \*\**P* < 0.0050).

To examine how microglia from cultured neocortical brain slices compare to cells *in vivo*, we leveraged the morphological reconstructions of microglia of primary somatosensory cortex of P15, P22, and P30-60 mice, randomly selected from the openly available database at neuromorpho.org ^41,42^. Considering that OSC slices are generated at P4/5, these *in vivo* time points covered the range before *ex vivo* experiments (P15, matching DIV9/10), while experiments were conducted (P22, matching DIV16/17), and when cells can be expected to be mature *in vivo* (P30-60, outside culturing range). We found that microglia *in vivo* at P22 showed the greatest similarity to parenchymal microglia in OSCs (**Figure 3E, F-H**). While slightly less ramified, P22 microglia did not significantly differ from parenchymal cells in quantified morphometric parameters, in contrast to microglia from P15, which tended to be slightly less complex. Interestingly, the morphology of parenchymal microglia appeared to be in an intermediate state between the age-matched P22 and the adolescent or early adult P30-60 time points (**Figure 3E** and **F**). These findings paint a diverse, yet broadly convergent picture of parenchymal microglia consistently displaying a ramified morphology, both within the OSC and when compared to age-matched microglia *in vivo*.

### Layer specific myelin maturation in cortical OSCs

Having found that microglia between DIV 14 and 21 are homeostatic, we next examined the time- and region-dependence of myelination in cortical OSCs. Staining with markers for myelin basic protein (MBP), axonal cytoskeletal marker neurofilament (NF) and paranodes with contactin-associated protein 1 (CASPR) at weekly intervals revealed three distinct phases of myelin development and stereotypic nodal organization of the axon-myelin units **(Figure 4A, B)**. Myelin coverage across the cortical lamina was determined by quantifying the area of myelinated and neurofilament-positive axons (MBP^+^ NFH^+^) divided by the total area of NFH^+^ axonal signal (myelination index, **Supplemental Figure 3**). Slices harvested at DIV 7 exhibited a sparse myelination index with a few oligodendrocytes in the deeper layers (**Figure 4A, C**). One week later there was a significant 8-fold increase in myelination across the entire cortex with stereotypic organization myelinated axons in superior layers and significantly higher degree of myelin coverage in the inferior layers which remained stable up to DIV 28 (**Figure 4A–C**). While axons within the white matter were heavily myelinated at DIV 14 (myelination index = 33.1 ± 15.5%) the region exhibited signs of myelin damage and swelling ^43^ **(Supplemental Figure 4),** consistent with the observed cell death and microglia activity in subcortical regions.

**Figure 4.**
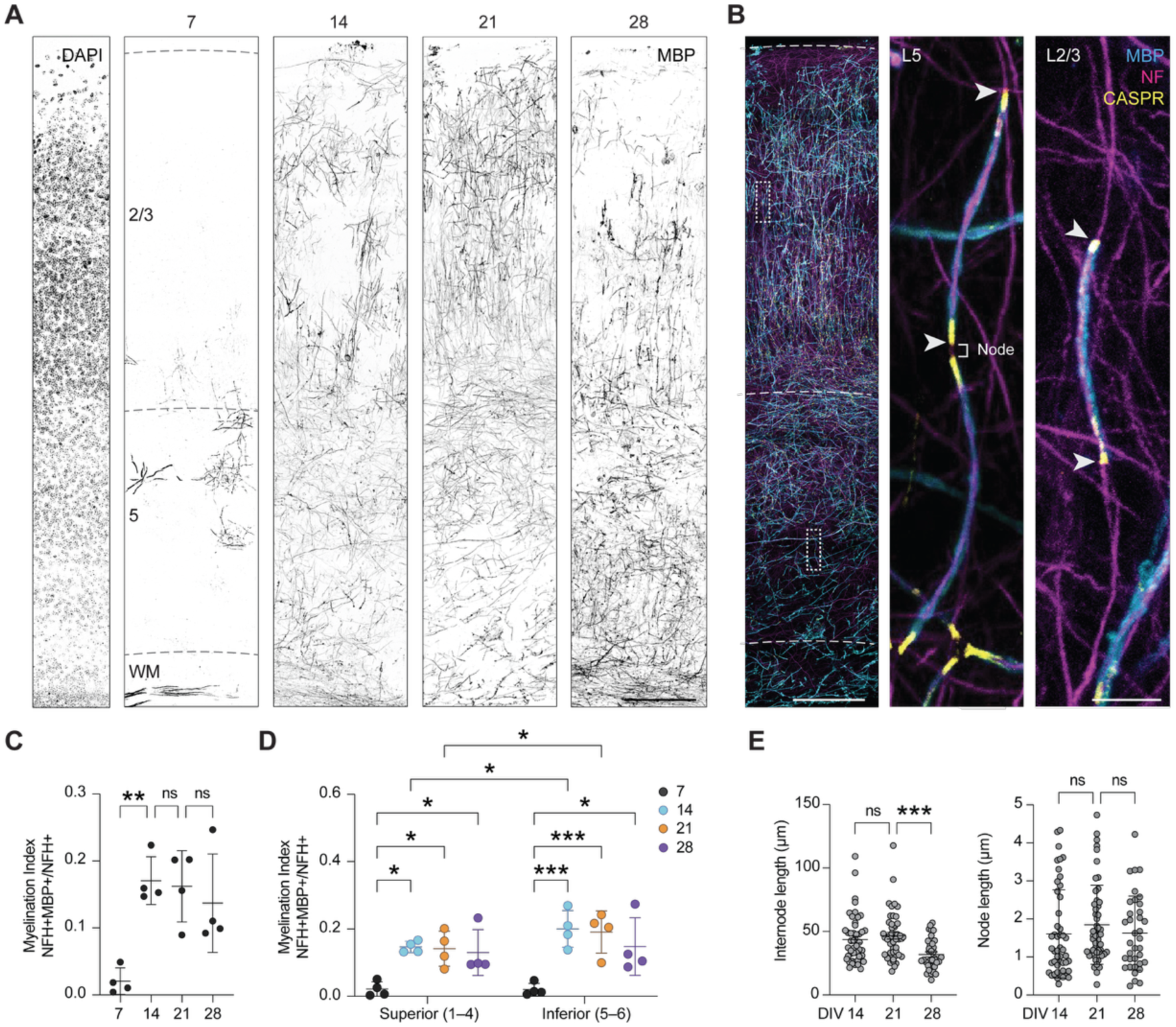
Myelin development in the OSC. **A.** Example *z*-projected confocal images of DAPI- and MBP-stained OSCs at 7-day intervals from DIV 7–28. Scale bar, 150 µm. **B.** (*Left*) Overview image of the combined MBP, NF and CASPR staining at DIV 14. Scale bar, 150 µm. (*Middle*) Confocal images of the nodal region and myelinated internodes from L5 and (*Right)* L2/3, from regions indicated in left image. Scale bar, 5 µm. **C.** Myelination index (percentage of NF area positive for MBP / total NFH positive area) increases from DIV 7–14 and is stable from DIV 14-28 (one-way ANOVA, ** *P* = 0.0051, *n* = 4 slices). **D.** Myelination index increases differentially between superior and inferior layers from DIV 7–14 and is stable from DIV 14-28. Inferior layers have increased myelination compared to superior layers from DIV 14-21, (two-way ANOVA, time *\*\* P* = 0.0034, region *\*\* P* = 0. 0065, interaction *P* = 0.1506, Tukey’s post hoc test *\* P =* < 0.05*, *** P =* < 0.001*, n* = 4 slices). **E.** (*Left*) Internode length is stable across DIV 14–21 but significantly decreases at DIV 28 (one-way ANOVA, *** *P* = 0.0004, *n* = 48 internodes for DIV 14/21 and *n* = 36 internodes for DIV 28). (*Right*). Node length is unchanged across DIV 14–28 (one-way ANOVA, *** *P* = 0.0004 *n* = 48 nodes at DIV 14, 51 nodes at DIV 21 and 36 nodes at DIV 28).

To investigate the layer dependent myelination of OSCs, we stratified the data by superior and inferior layers. Analysis of myelin coverage within regions and across development revealed significant changes in myelination index were driven by both time and region (Two-way ANOVA; time *\*\*P* = 0.0034, region *\*\*P* = 0.0065, interaction *P* = 0.1506). Both superior and inferior cortical layers exhibited a significant increase in myelination index from DIV 7–14 (7- and 10-fold increase, respectively), and myelin coverage remained stable from DIV 14 to 28 (**Figure 4D**).

Interestingly, consistent with the patchy myelination of pyramidal neuron axons in the superior layers ^44^ (**Figure 4B**), the superior cortical regions showed a significantly lower myelination index, in comparison to inferior regions suggesting that the differential myelin patterns, with inside-out gradients, develops intrinsically (**Figure 4D**). We further examined myelin development at the level of single axons and quantified the internode length (distance between two CASPR puncta along a MBP+ axon) and as a proxy of nodal length measured the distance between two flanking CASPR puncta in inferior layers from DIV 14, 21 and 28 (**Figure 4A–D**). The results showed that while the node length remained stable, on average ∼1.5 µm, internode length significantly decreased in the fourth week consistent with a decreased myelin integrity (**Figure 4E**). Together, these findings indicate that myelin development in culture resembles the in vivo organization across cortical layers. The timing of myelin coverage also coincides with microglial homeostasis (**Figure 2, 3**) and further implicates the DIV 14-21 period as an optimal time point for glial cell investigations.

### Distinct features of myelin ultrastructural composition in OSCs

The MBP staining suggests the presence of compacted myelin sheaths. Previous slice cultures studies using transmission electron microscopy (TEM) revealed compact myelin after several weeks ^45–47^. To examine the ultrastructural features of myelin in the OSCs the slice cultures on DIV 21 were fixed, stained for TEM and imaged at >20,000 magnification. Myelinated axons showed stereotypic organization at the ultrastructure. **Figure 5A** shows a transverse section of an axon containing in the axoplasm numerous neurofilaments and microtubules, being wrapped by a myelin sheath, with compact myelin as well as inner- and outer tongues and radial components. Being that OSCs were prepared from P4-5 pups, DIV 21 slices are thus 25-26 days old. Based on 89 cross-sectional images of myelinated axons we measured myelin thickness across a range of axon calibers and compared the results with TEM from *in vivo* somatosensory cortex of a C57BL/6J mouse of similar age (**Figure 5B**).

**Figure 5.**
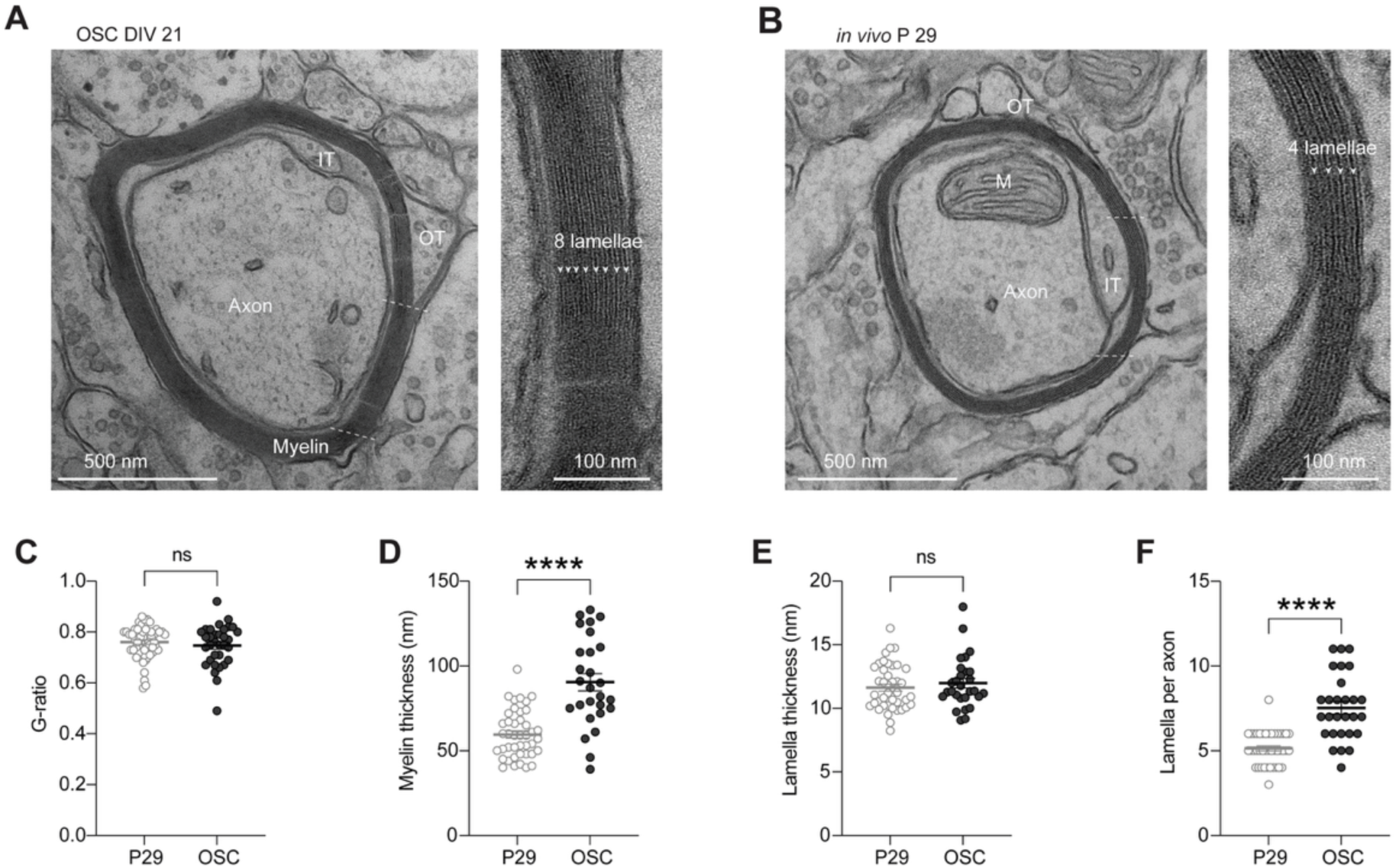
Myelin ultrastructure in the OSC. **A.** Left, representative TEM image of an axon from a DIV 21 OSC (axon diameter 851 nm, myelin thickness 96 nm) with stereotypic ultrastructure of compact and noncompact myelin. The inner tongue (IT) and outer tongue (OT) of the oligodendrocyte are indicated in white text. Scale bar 500 nm. Right, zoom in showing the layering of 8 lamellae. Scale bar 100 nm. **B.** Left, representative TEM image of an axon from an *in vivo* preparation P29. Axon diameter 655 nm, myelin thickness 41 nm, showing compact and noncompact myelin. The inner tongue (IT) and outer tongue (OT) of the oligodendrocyte and axonal mitochondria (M) are indicated in white text. Scale bar 500 nm. Right, zoom in showing the layering of 4 lamellae. Scale bar 100 nm **C.** OSC axons exhibit a trend toward lower g-ratio compared to age matched axons from *in vivo* preparations (Mann-Whitney test, *P* = 0.0502, *in vivo n* = 56, OSC *n* = 33). **D.** OSC axons have thicker myelin compared to age- and diameter matched axons from *in vivo* preparations (Mann-Whitney test, \*\*\*\**P* = <0.0001, *in vivo n* = 42, OSC *n* = 28). **E.** Individual lamella thickness in OSCs is similar to lamella thickness in age- and diameter matched axons from *in vivo* preparations (Mann-Whitney test, *P* = 0.5064, *in vivo n* = 42, OSC *n* = 28). **F.** OSC axons have more lamellae compared to age- and diameter matched *in vivo* axons (Mann-Whitney test, \*\*\*\**P* = <0.0001, *in vivo n* = 42, OSC *n* = 28).

While axons analyzed from cortical OSC and the *in vivo* brain had mean axonal diameters, 0.78 µm and 0.70 µm respectively, axons greater than 1 µm or smaller than 0.5 µm were absent in the OSC (**Figure 5C)**. We then compared *g*-ratio of axons across a comparable range of axonal calibers (0.5–1 µm). These data showed that OSC axons had a trend toward lower g-ratios (from 0.7602 to 0.7473, *P* = 0.0502, **Figure 5D**). To further explore possible differences underlying the OSC myelin ultrastructure we measured the sheath thickness and found that cortical OSCs had significantly thicker myelin sheaths compared to age-matched in vivo preparations (**Figure 5E**). Furthermore, by counting single myelin lamellae, we discovered that the sheaths were composed of lamella with similar thickness (∼12 nm), but significantly more myelin membranes within one sheath (**Figure 5F, G**). Thus, OSCs are characterized by topographically normal neocortical myeloarchitecture with relatively thicker myelin sheaths.

### Neuronal network in OSCs intrinsically generate UP-state activity

Microglial states and myelination are known to fbe influenced by neuronal activity ^2,26,46,48^. To determine the specific electrophysiological activity patterns of neurons in the OSC we made whole-cell patch clamp recordings from L5 pyramidal neurons identified by the *Rbp4*-Cre driver line, or using visual identification of neurons in slices from *Hexb*^tdTom^ slices. The electrophysiological properties were assessed at DIV 14–21 and in some cases the morphology was examined by filling cells with biocytin (n = 5, **Figure 6A**). The resting membrane potential was on average –74.8 ± 1.3 mV, with an input resistance of 90.2 ± 10.6 MΩ (n = 13). By injecting steady-state depolarizing currents we observed that the pyramidal neurons exhibited firing properties akin to regular-firing type L5 pyramidal neurons, with firing frequencies of 13.7 ± 1.8 Hz at rheobase (**Figure 6A-C**). Remarkably, burst-firing L5 pyramidal neurons, characterized by a cluster of multiple action potentials with an inter-spike interval < 10 ms, a hallmark of the thick-tufted subcortically projecting L5 pyramidal neurons ^49^, was not observed (0 out of 13 cells).

**Figure 6.**
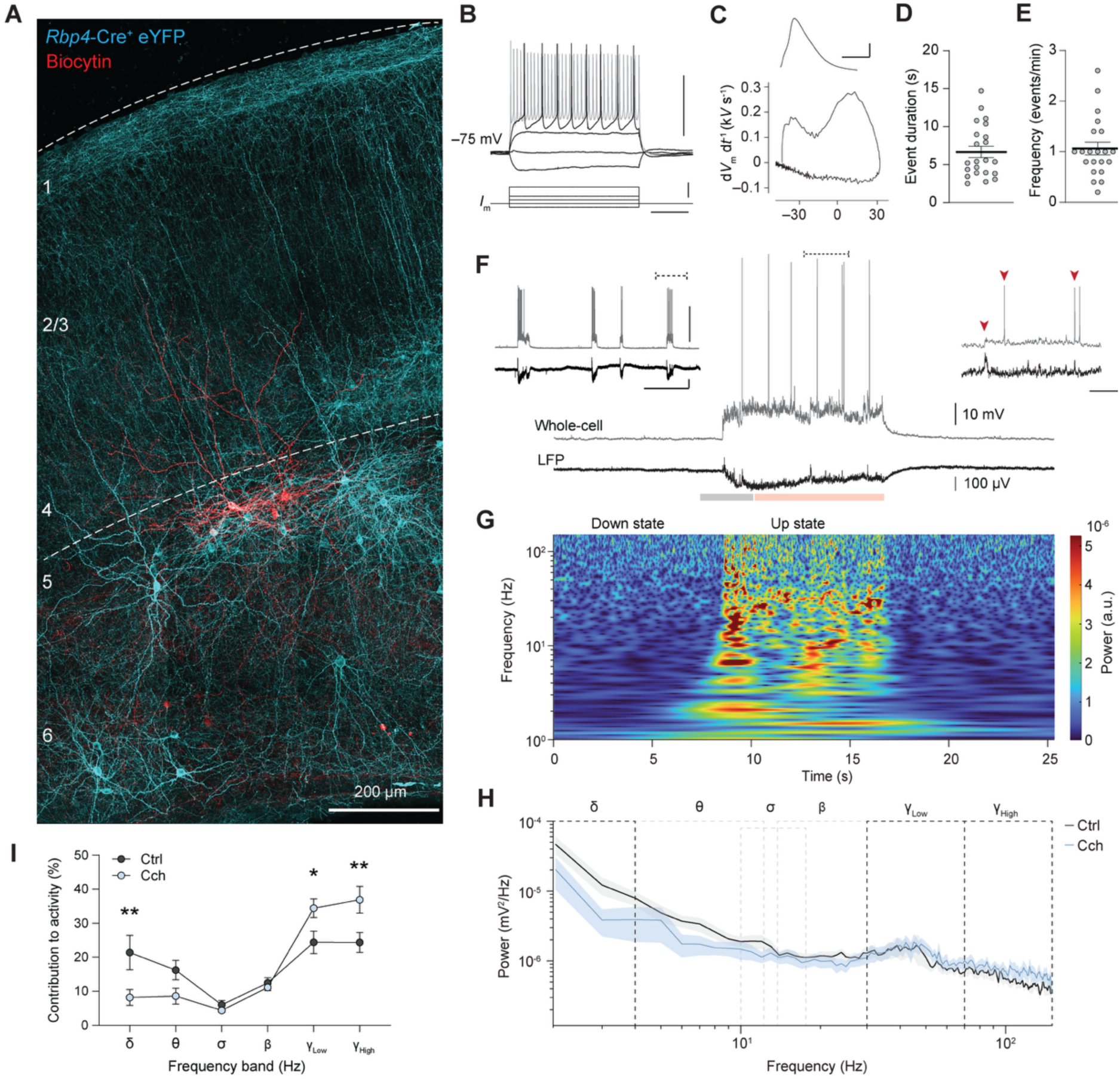
Neuronal Activity Patterns in OSCs. A. Z-projected confocal image of a *Rbp4*-Cre OSC (DIV 21) transfected with AAV5-ChR2-eYFP. Staining for eYFP (cyan), patch-clamp and biocytin-filled pyramidal neurons (red). B. Family of voltage responses to step-current injections of a L5 pyramidal neuron. Scale bars, 50 mV, 100 pA and 100 ms. C. Phase-plane plot of the action potential. Inset shows the voltage-time of the action potential, scale bars, 20 mV and 1 ms. D. Population data of Up-state duration. Dots indicate averages from a 5 minute time window (n = 22 slices), thick bar indicates mean and error bars SEM. E. Population data of Up-state frequency. Replicates and bars as in D. F. Example Up-state from a pyramidal neuron with combined whole-cell recording (light grey) and LFP recording (dark grey). Baseline segment of the Up-state in light grey, onset period in grey, and Up-state body in orange. Inset to the left shows minute time-scale event occurrences (time axis 1 minute), with the dotted line indicating main trace location, while the dotted line on the main trace indicates the location of the inset to the right, showing millisecond time scale (time axis 500 ms) to highlight subthreshold depolarizations and variable event-locking between LFP and whole-cell, which itself is indicated with red triangles. G. Time-frequency diagram showing the frequency distribution of neuronal network activity during the Down- and Up-state, taken from the LFP trace in F. H. Power spectral density diagram of LFP activity from slices in control (Ctrl) and carbachol-treated (Cch) condition. I. Carbachol induces gamma oscillations at the expense of delta activity. Percentage of band-integrated power over total power for indicated frequency bands in control (Ctrl) and carbachol (Cch) condition (two-way ANOVA, frequency band \*\**\*\* P* < 0.0001, treatment *P* = 0. 0.8838, slice *P* = 0.1021 interaction **** *P* < 0.0001, Tukey’s post hoc test *\* P =* < 0.05*, ** P =* < 0.005*, n* = 6 slices).

Cortical slices are known to generate large synchronous population bursts – so-called Up-states – followed long silent periods, which are shaped by recurrent connectivity, synaptic inputs, and intrinsic membrane conductances ^25,27,50^. To record these events, we paired local field potentials (LFP) with the whole-cell recordings of pyramidal neurons. Slices of 14–21 DIV exhibited Up-state activity at frequencies of approximately 1 event/minute, and a duration of multiple seconds (on average 6.7 ± 0.7 s, n = 22 slices, **Figure 6D-F**), in line with previous reports ^25^. Studying the Up-states in more detail, we found that LFP and whole-cell recordings generally showed a strict coherence between the intracellularly recorded membrane potential and network-level voltage shifts that delineate Up-state episodes (**Figure 6F**). To examine the frequency composition and temporal dynamics of Up-states, we generated a time-frequency plot which showed a broad diversity of activity patterns within the events (**Figure 6G**). Within the steady state of the Up-state, large and heterogeneous oscillations could be observed that overall showed a peak in the low gamma range (30–60 Hz, **Figure 6H**), suggesting a key contribution of fast-spiking parvalbumin interneurons. To test their contribution to the network oscillations, we used the cholinergic agonist carbachol, known to impact *in vivo* Up-states and acting predominantly on fast-firing PV^+^ basket cells ^51^ . Bath application of 20 µM carbachol showed that Up-state spectral properties shifted significantly, with low-frequency activity significantly decreasing in the delta range, while high frequency firing in both low and high gamma bands increased (**Figure 6 H, I**). Taken together, neocortical OSCs display electrophysiological properties comparable to cortical tissue *ex vivo*. Intrinsically generated Up-state activity in OSCs harbors stereotyped frequency components that can be altered via neuromodulator manipulation, enabling the study neuroglial responses to alterations in neuronal firing activity.

### Oligodendrocyte morphology and glia-glial interactions in live tissue

The results indicate that the DIV 14–21 time point recapitulates key aspects of homeostatic glial and neuronal function. Shortly before this period microglia transition into a homeostatic phase while oligodendrocytes undergo a wave of myelination. Given the prominent network activity, we explored the possibilities to perform live imaging. To label oligodendrocytes in OSCs we first used AAV2/1-*Mbp*:mem-EGFP with a membrane bound fluorophore ^32^ and verified the selective labeling by post hoc staining with a MBP antibody (**Figure 7A**). Colocalization analysis revealed 73.8% of all EGFP signal to be colocalized with MBP antibody labeling. To assess whether the EGFP labeled structures indicated the presence of myelin, EGFP positive internodes were then individually segmented and colocalization analysis revealed them to be nearly entirely covered with MBP (**Figure 7B**). Use of the membrane bound EGFP signal allowed for the visualization of both MBP positive myelin sheaths and cytoplasmic filled process leading back to oligodendrocyte soma, and the imaging resolution with live 2-photon microscopy allowed to distinguish smaller membrane extensions including paranodal bridges ^52^ (**Figure 7A, C) (Supplemental Figure 5)**.

**Figure 7.**
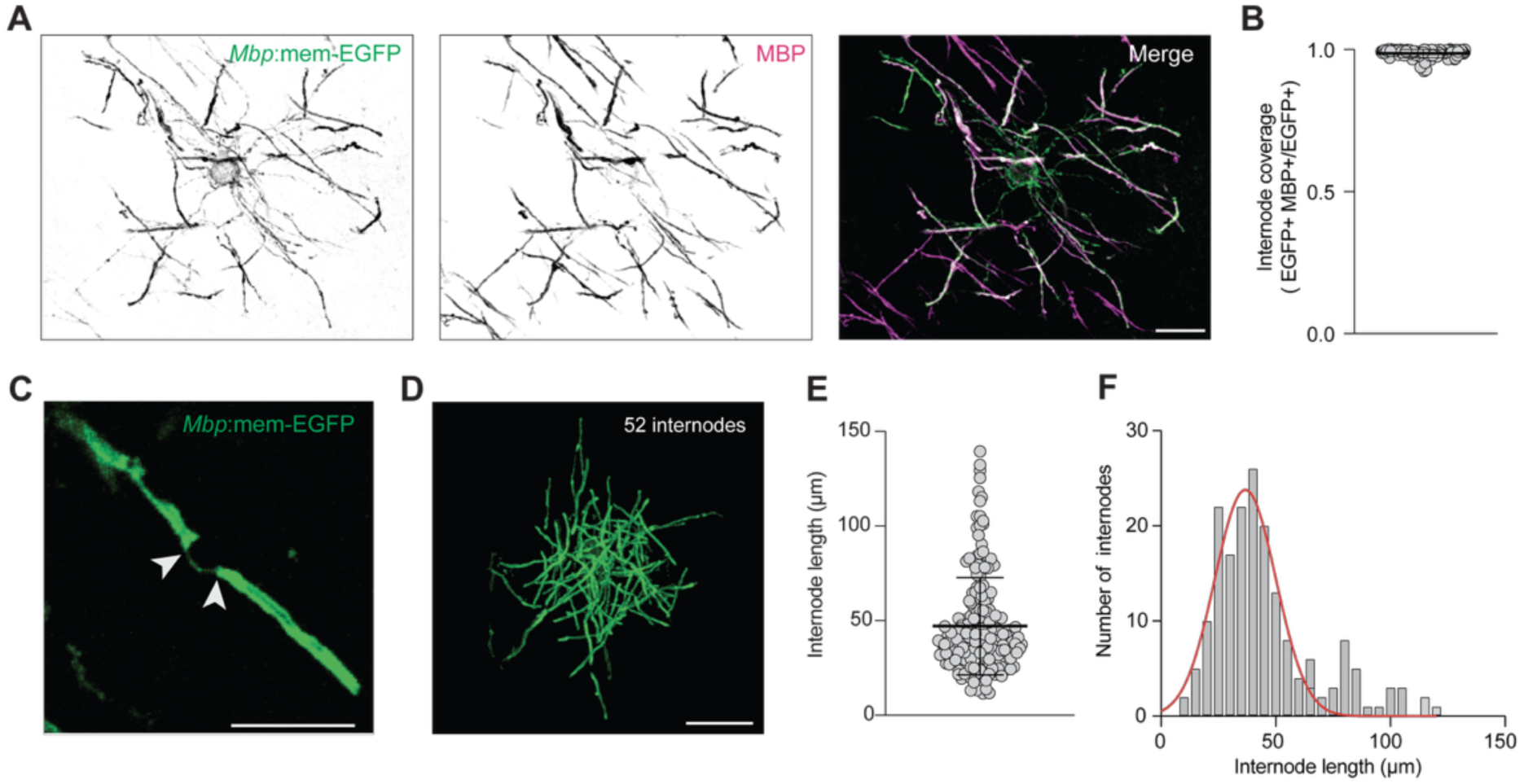
Oligodendrocyte morphology in OSCs. **A.** Representative confocal image of an oligodendrocyte from DIV 21 OSC transfected with AAV2/1-*Mbp*:mem-EGFP and stained for MBP. Mem-EGFP (left, green), MBP antibody (middle, magenta), merge (right). Scale bar 20 µm. **B.** Fluorescence colocalization of MBP and EGFP labeled internodes yielding high overlap, 98.6 ± 1.48 %, *n* = 99 internodes. **C.** Z-projected live two photon image of myelin sheaths linked by paranodal bridge indicated by arrows. Scale bar 20 µm. **D.** Z-projected live two photon image of a single oligodendrocyte with 52 internodes. Scale bar 40 µm. **E.** Population data of internode length. Mean internode length of 47.13 ± 25.63 µm, *n* = 188 internodes, N = 5 oligodendrocytes. **F.** Histogram of internode length. Data fit with Gaussian function (R^2^ = 0.8659), *n* = 188 internodes, from 5 oligodendrocytes.

The sparse labeling of oligodendrocytes enabled us to quantify the specific number and length of internodes per cell in cortical OCSs. Careful tracing of myelin sheaths revealed each oligodendrocyte had ∼38 ± 10 internodes per oligodendrocytes (n = 5, **Figure 7D**) with an average internode length of 47.13 ± 25.63 µm (n = 188, **Figure 7D, E**). Both values are in alignment with metrics previously reported for internode morphology in L5 of the mouse somatosensory cortex under *in vivo* conditions ^53^. Together, these measurements indicate the viral labelling strategy is a powerful tool for studying oligodendrocytes live in the highly manipulatable environment of cortical OSCs.

Having verified successful oligodendrocyte labeling we next characterized microglia-myelin interactions in cortical OSCs. These glia-glia interactions are implicated in the process of, myelin development, maintenance, removal of damaged sheaths ^9,54^, and these key functions involve dynamic microglial process movements within the parenchyma ^26,54,55^. To better understand the duration of transient microglia myelin interactions we utilized the *Hexb^tdTom^* reporter mouse line in combination with AAV2/1-*Mbp*:mem-EGFP. We employed live 2-photon imaging to track microglial surveillance of myelin sheaths in culture over brief 30 min imaging sessions (**Figure 8A**) and observed ramified motile microglia extending thin processes towards myelin sheaths and making direct contact (**Figure 8B–D**). This contact was then followed by microglial process retraction as the cells continued to survey the surrounding space. To visualize the transient nature of microglial cell process extensions and retraction we color coded individual frames of the time series (**Figure 8E**). The dynamic color range of microglia processes and solid white color of myelin sheaths and microglia cell bodies indicate the soma of the microglia remained stationary during homeostatic surveillance. Intensity profiles of microglial fluorescence (tdTom) and myelin sheaths (EGFP) were gathered from microglia-myelin contact sites, and the peak microglia fluorescence were aligned to generate a normalized timeline of transient contacts (**Figure 8F**). The average duration of contact was found to be ∼10 min (**Figure 8G**). Previous work in the neocortical OSC showed microglia take up multiple myelin lamella, forming myelin bodies, pulled away from internodes via ‘myelin tethers’ ^56^. Consistent with myelin tethers, which are rare to capture live, post-hoc staining with Iba1 and MBP and full field examination showed that consistently these myelin pruning events contained microglia engulfed myelin (**Figure 8H**).

**Figure 8.**
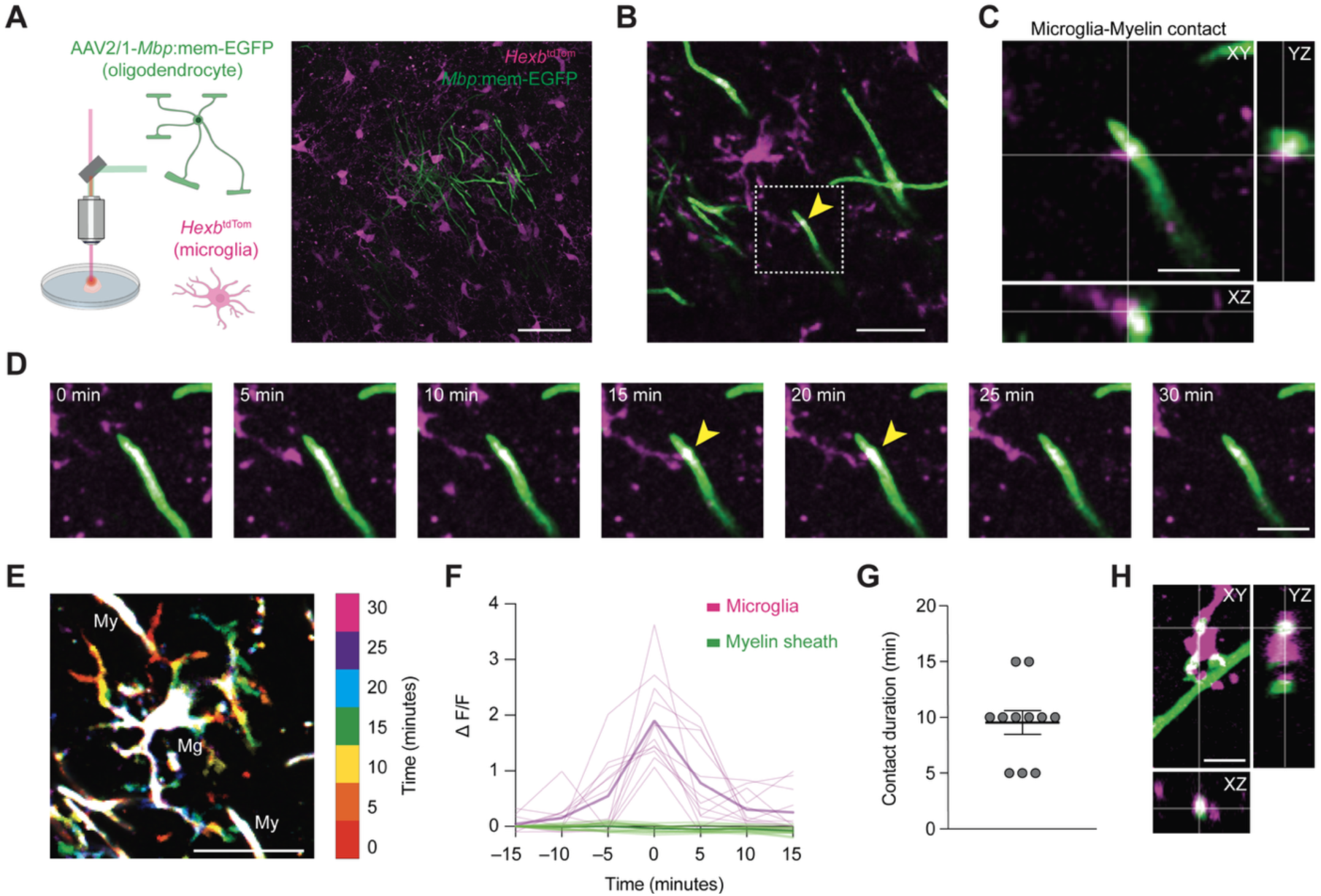
Baseline neuroglia-interactions. **A.** Schematic of 2-photon imaging visualizing microglia (*Hexb*^tdTom^, magenta) and oligodendrocytes (AAV2/1-*Mbp*:mem-EGFP, green). *Z*-projected live 2-photon image examples. Scale bar, 50 µm. **B.** *Z*-projected live 2-photon image of single microglia contacting a myelin sheath. White box and yellow arrow indicate the interaction site. Scale bar, 20 µm. **C.** Orthogonal view of the microglia-myelin contact site from the inset in B. Scale bar, 10 µm. **D.** Images of microglia myelin interaction at 5 min intervals. Yellow arrow indicates contact point. Scale bar, 10 µm. **E.** Image of microglia and oligodendrocytes color coded by time. Microglia and myelin sheaths are indicated by Mg and My respectively. Scale bar 20 µm. **F.** Intensity profiles of microglial contacts with myelin sheaths aligned to the peak fluorescence during contact. (*n* = 11 microglia myelin contact sites.) **G.** Quantification of contact duration (*n* = 11 microglia myelin contact sites.) **H.** Orthogonal view of a post-hoc confocal image of microglia (Iba1, magenta) with engulfed myelin tether (MBP, green). Scale bar, 5 µm.

## Discussion

By studying neurons and glial cells in an integrated manner we show that neocortical OSCs exhibit molecular and structural features of homeostatic cortical tissue, including retained cortical layering and glial cell development. Myelinating oligodendrocytes and homeostatic, highly ramified microglia arise and are maintained in three distinct phases, following a concerted developmental maturation that culminates in molecular and morphological properties similar to those found *in vivo*.

As the main phagocyte of the brain, we find, as expected, that microglia become activated upon slice preparation. However, after two weeks in culture the cells take up a homeostatic profile through P2Y12R expression as well as a highly ramified morphology (**Figures 2, 3**). This development contrasts with microglia *in vivo* showing that P2Y12 can be readily detected with increasing expression from P6 onwards ^57^. The delay in microglial expression of P2Y12R in OSCs is likely explained by microglia reacting to slicing, whereby they dramatically change their cellular state, transitioning from surveillant cells to being primarily phagocytically active ^58^. Interestingly, their ability to still take up homeostatic profiles contrasts with data showing that insults like traumatic brain injury ^59^, or immunological challenges ^60^, can leave microglia in a primed state unable to return to homeostasis. It is thus likely that the microglial state in culture is neither purely subject to developmental plans nor to environmental stimuli but instead reflects a product of the two.

The observed sustained CD68 expression suggests continuously ongoing phagocytosis across the life span of the organotypic culture. What microglia are phagocytosing is not well understood and remains to be examined in future studies. It is likely microglial phagocytosis targets synaptic structures as part of the network maturation and/or myelin sheaths which rapidly develop in this phase. Importantly, the P2Y12^High^/CD68^High^ state is not a common expression combination fitting to a known cluster, underscoring the idea that microglia instead exist along a continuum of states that – up to a point – can give way to one another ^61^. Previous studies have shown that ramification at a homeostatic-like state can be found in microglia in a neocortical slice culture context despite concomitant CD68 expression ^62^, and transcriptomic analyses of OSC microglia, CD68 expression remained higher than age-matched acutely isolated cells, even after weeks in culture ^21^. The presence of ramified microglia in parenchyma of organotypic slices is in line with morphological studies in slice cultures ^63–65^ ^20,62,66^. However, the present findings advance these studies by providing spatiotemporal insight into the molecular signature of microglia along the cortical column, as well as the morphology of single microglia at various locations within a cultured slice.

Another key finding of the present research is that the microglial state changes do not happen in isolation but instead reflect a general development and maturation of the slice, as can be seen by myelination following a similar tri-phasic developmental epoch. The first stage was characterized by sparse myelin coverage (DIV 0–7). This window of slice development is also home to ameboid microglia and low levels of P2Y12 expression. While our observed delay in P2Y12 expression may be influenced by slice related debris clearance, previous work *in vivo* has described a necessary phase of microglia-oligodendrocyte linage cell interactions whereby ameboid phagocytically active cells are essential in establishing developmental myelination^10,67,68^ . Microglia during this time lack classic “homeostatic” markers of adult microglia but are nonetheless important for creating a permissive environment for OPC differentiation via clearance of whole cells and release of promyelinating factors^67^ . The adult oligodendrocytes emerging in the second stage (DIV14–21) produced compact myelin sheaths and a laminar-dependent myeloarchitecture recapitulating patterns observed *in vivo*, with sparse myelination in superficial layers and denser and more continuous myelination of the inferior layers ^44,53^. OSCs maintain an innate capacity to myelinate due to oligodendrocyte precursor cell (OPC) colonization by E16 (ref. ^69^), which further differentiate to produce mature myelinating oligodendrocytes ^70^. From DIV 21 onward we observed a spontaneous region-specific cell death occur in deeper layers. This wave of death proceeds overt changes in slice health observed at DIV 28 where we find increased variability in both microglial and oligodendrocyte markers in combination with shortening of internodes. Together, this paints a picture of active slice degeneration from DIV 21–28 and this timepoint may therefore be leveraged by future studies to interrogate the neuro-glia interactions responsible for slice culture decay. Of particular interests would be to uncover the contents of the lysosomal compartments which remain in abundance across the culturing window.

Remarkably, we found that axons in the OSC had more myelin wraps when compared to age-matched *in vivo* axons (**Figure 7**). What explains this increased myelin thickness is not well understood. One possibility is that the deeper layers of the OSC, are composed of a distinct ratio of inhibitory to excitatory axons, which are known to differ in myelin composition at the ultrastructure and molecular level ^71–73^. To test this possibility, future studies require using immunogold EM or other cell-type specific labeling methods to identify the myelinated axons in the OSC. Another possibility is an impaired astroglia or microglia myelin maintenance leads to an axonal hypermyelination ^9^. Microglia are able to selectively prune individual lamella away from the underlying sheaths without effecting the underlying myelin integrity ^56^ or participate in whole sheath phagocytosis ^74^. This role could be tested by depleting the microglia or inhibiting microglial uptake of established myelin to uncover mechanisms of myelin maintenance. The organotypic culture offers an excellent platform to test such key causality relationships, and time-lapse imaging of cell types over days would substantially further strengthen evidence.

An alternative and intriguing explanation for the increased myelin thickness is that the slowly oscillating highly synchronized bursts of neuronal activity in the OSC may promote myelin wrapping. Corticothalamic circuits generate highly rhythmic Up-state depolarizations which are key rhythms for the coordination of synaptic plasticity, and behavioral wake- and sleep states including the non-rapid eye movement sleep ^75^ . These oscillations originate in the cortex since they continue even when the cortex is disconnected from thalamic input and they can be observed in cortical slices, suggesting they emerge intrinsically from synaptically connected neocortical neurons ^76^. The stereotypic Up-state events we observed (**Figure 6**) showed similar duration and frequency as described previously in slice cultures of the neocortex ^25,27^. In addition, our spectral decomposition of the LFP revealed a broad range of band power, peaking around 20–60 Hz, closely corresponding to gamma activity. This is further in line with previous studies showing that PV interneuron activity in this band sharply increases around DIV 14, and their rapid perisomatic inhibition coordinates gamma activity and Up-states ^25,77,78^. Unlike in vivo conditions, however, the Up-states in cortical slices are persistent and represent the only network mode. Whether the rhythmic slow oscillations, through mechanisms of activity-dependent myelination ^48,79^, facilitate myelogenesis needs to be further investigated. Blocking voltage-gated channels or using neuromodulators directly regulating neuronal excitability could be leveraged to modulate network events and dissect the molecular- and activity-dependent mechanisms of cell-type specific myelination.

Taken together, our live imaging and cellular characterization highlights the spatial and temporal resolution of this platform for visualizing neuron-microglia-myelin interactions. Future studies may use the window from DIV14–21 to better understand how microglia mediate myelin maintenance. Neocortical tissue in the organotypic culture thus presents a valid and flexible platform for mechanistic studies of the molecular and cellular basis of neuroglial interactions in health and disease.

## Methods

### Animals

All animal procedures were done in after approval from the Royal Netherlands Academy of Arts and Sciences (KNAW) Animal Ethics Committee (DEC) and Central Authority for Scientific Procedures on Animals (CCD, license AVD80100202216329). The specific experimental designs with animals were evaluated and monitored by the Animal Welfare Body (IvD, protocols NIN.21.21.01, NIN.21.21.07, NIN.22.21.02). For wildtype mice we used male and female C57BL/6 mice. To visualize microglia during live imaging experiments we used mice expressing tdTomato under the *Hexb* promoter ^30^ and prepared slice cultures from homozygous (*Hexb*^tdTomato/tdTomato^) animals. Mice were acquired from the university of Freiburg and obtained from a new colony at the NIN. The *Rbp4-Cre* mouse strain used for this research project was B6.FVB(Cg)-Tg(*Rbp4*-cre)KL100Gsat/Mmucd, RRID:MMRRC_037128-UCD, obtained from the Mutant Mouse Resource and Research Center (MMRRC) at University of California at Davis, an NIH-funded strain repository, and was donated to the MMRRC by MMRRC at University of California, Davis. The strain was made from the original strain MMRRC:032115 donated by Nathaniel Heintz, Ph.D., The Rockefeller University, GENSAT and Charles Gerfen, Ph.D., National Institutes of Health, National Institute of Mental Health.

### Cortical organotypic slice culture preparation and culturing

Cortical organotypic slice cultures were prepared from 4-5 day old mouse pups. Briefly, mice were anaesthetized via hypothermia, then sacrificed by decapitation with scissors. The brain was extracted and placed in ice-cold dissection solution consisting of 98% GBSS (in mM) 1.5 CaCl_2_, 0.2 KH_2_PO_4_, 0.3 MgSO_4_, 2.7 NaHCO_3_, 5.0 KCl, 1.0 MgCl_2_, 137 NaCl, 0.85 Na_2_HPO_4_, 5.6 D-glucose), 1% (0.1 M stock) kynurenic acid, and 1% (2.5 M stock glucose), 1% (0.1 M stock kynurenic acid), sterile filtered with 0.2 µm filtration flasks, and adjusted to pH 7.2 and 320 mOsm. Under a dissection microscope, brains were cut down the midline with a scalpel and then sectioned via McIlwain Tissue Chopper to obtain 300 µm thick coronal slices. Slices were quickly transferred to hydrophilic PTFE membrane inserts (Merck-Millipore, PICMORG50) in 6-well plates and cultured at 35 °C and 5% CO_2_ in culturing medium consisting of 47.75% MEM (Thermo Fisher Scientific # 11575032), 25% HBSS (Thermo Fisher Scientific # 24020133), 25% heat-inactivated horse serum (Thermo Fisher Scientific # 26050088), 2% (2.5 M stock) D-glucose, and 1.25% (1 M stock) HEPES (Sigma-Aldrich H3375), sterile filtered, and adjusted to pH 7.2 and 320 mOsm. Medium was changed 3 x a week with fresh equilibrated medium. AAV2/1-*Mbp*:mem-EGFP or AAV5/Ef1a-DIO-hChR2(E123T/T159C)-EYFP (RRID:Addgene_35509) was applied directly to slices cultures at DIV 7 to selectively label oligodendrocytes and L5 pyramidal cells.

### Immunohistochemistry

Cortical organotypic slice cultures were removed from culturing medium and placed directly in 4% PFA in PBS for 20 min followed by 3 washes in PBS for 10 min. The slices were blocked for 2 hrs in PBS containing 0.3-1% Triton X-100 and 10% normal goat serum. Next, slices were incubated in buffer containing half the Triton and goat serum concentrations in PBS, as well as primary antibodies (Supplemental Table 1), overnight at room temperature while shaking. After primary incubation, slices were washed 3 times for 10 min in PBS. Following washing steps, slices were incubated in PBS containing secondary antibodies (Supplemental Table 1), as well as Triton and goat serum concentrations equal to those used for primary incubation, for 2 hrs at room temperature while shaking and protected from light. The slices were given final washing in PBS (3 x 10 min) before mounting with Vectashield mounting medium containing DAPI.

### Propidium iodide live/dead assay

Propidium iodide was added to equilibrated medium at 5 µg/ml final concentration and the complete medium of cultures was exchanged. To ensure penetration of the solution on the complete surface area of the slice, a few droplets of media were gently placed onto the slices, with run-off being aspirated from the membrane. Slices were incubated for 1 hr in the incubator before being washed twice with pre-equilibrated PBS. Subsequently, slices were fixed in 4% PFA in PBS for 20 minutes while being protected from light and thereafter washed 3 times for 10 minutes in PBS. As a counterstain, slices were incubated in DAPI (1:1000, find final conc) in PBS for 20 minutes and washed 2 times in PBS. They were then mounted in Vectashield with DAPI.

### Confocal and apotome imaging

Immunohistochemically stained slices were imaged on a Leica Microsystems SP8 confocal laser scanning microscope using a 63x 1.4 NA or 40x 1.3 NA lens and running LAS X v.3.5.7. Tile scans of cortical columns were generated using sequential scans of individual channels and imported to FIJI for colocalization analysis with BIOP JACoP. Axons within inferior layers were manually selected and internode and nodal distance were measured along MBP^+^ signal or between two CASPR positive puncta respectively using the SNT plugin ^80^. Colocalization analysis of virally labelled oligodendrocyte with MBP^+^ signal was achieved by first segmenting individual EGFP positive internodes in SNT before running BIOP JACoP. For microglial state analysis, each channel was thresholded and de-noised using binary operations and punctal size exclusion, then colocalization between channels was assessed via the image calculator, measuring the area fraction positive for each respective channel. Propidium iodide stained slices were imaged on a Carl Zeiss Apotome 3 optical sectioning microscope with a 20x 0.8 NA objective lens and Zeiss ZEN Blue v3.7.

### Live imaging and cell tracing

Two-three-week old cultures were cut so that individual, membrane-attached brain slices could be transferred from well plates to the bath of a two-photon microscope (Femto-2D-RD, Femtonics), perfused with carbogen-bubbled recording ACSF warmed to 30–33 °C. *Hexb*^TdTom^ labelled microglia or mem-EGFP labelled oligodendrocytes in combination with *Hexb*^TdTom^ labelled microglia were visualized via a Ti:Sapphire laser (Chameleon Ultra II; Coherent, Inc.) tuned to 1030 nm and 980 nm respectively, and cells were imaged using a 1.0 NA 60x or 1.0 NA 20x lens (Olympus) with pixel sizes between 0.2 x 0.2 x 1 µm (X/Y/Z) to visualize microglia and 0.3 x 0.3 x 1 µm to visualize oligodendrocytes in combination with microglia. For microglial morphometric data, imaged volumes spanned the entire depth of the slice culture (roughly 120-180 µm in *z*-distance). Image acquisition was performed with the MES software v6.3.7902 (Femtonics). Fields of view were selected to maximize the *z*-depth in the cortex, roughly in the inferior L2/3 for microglial reconstructions and L5 for oligodendrocyte containing images. From acquired micrographs, individual cells were selected based on the intactness of their cellular arborizations, traced, and analyzed using the SNT plugin for FIJI ^80,81^. Openly available morphological reconstructions were randomly selected from the Siegert dataset on the Neuromorpho platform and analyzed as described above (NeuroMorpho.Org, RRID:SCR_002145^41,42^ . Microglia myelin contact duration was determined by selecting regions of interest with a microglia-myelin contact and calculating ΔF/F across the imaging sessions. The maximum peak of each recording was then aligned and to create a normalized timeline of contact duration.

### Transmission Electron Microscopy

Insert with attached OSC was fixed for 1 hour with freshly made 4% paraformaldehyde (Sigma-Aldrich, prilled) + 2.5% glutaraldehyde (aqueous solutions, EMS) in OSC medium. Afterwards, the insert with OSC was transferred to fresh fixative in 0.15M cacodylate buffer (aqueous solution, EMS) and stored at 4 °C for 48 hours. Samples were cut out of insert and removed from mesh, washed in 0.15M cacodylate buffer (aqueous solution, EMS) and postfixed with 1% osmium tetroxide (aqueous solution, EMS), 1.5% potassium ferrocyanide (EMS) in 0.15M cacodylate buffer (EMS) for 45 minutes. Samples were washed with MilliQ and dehydrated with increasing concentrations of EtoH; twice with 50% and once in 70%, 80%, 90% for 15 min each, and twice in 100% EtoH for 20 min each. Samples were penetrated with 1:1 EtoH-resin mixture for 1 hour and another hour with 1:3 EtoH-resin mixture. Samples were placed in fresh resin (Embed 812 EMS) and stored for 24 hours at 37 °C and polymerized at 65 °C for 72 hours. Samples were cut with a diamond knife (Diatome) on a Leica ultracut UCT and placed on formvar covered copper grids (EMS). Sections were post-stained with uranyl acetate (3.5% in ddH2O) for 20 min and lead citrate (EMS) for 2 min. For the *in vivo* EM a P29 BL/6 wildtype mouse was perfused with 20 mL 0.1M phosphate buffer followed by 20 mL 3% paraformaldehyde and 3% glutaraldehyde (EMS) in 0.1M phosphate buffer. Thick vibratome sections (100 µm) were post-fixed in similar fixative for 1 hour. Sections were transferred to storage buffer of 0.5% paraformaldehyde in 0.1M phosphate buffer. A region of interest was cut out from the slice and prepared for resin embedding as described above.

All samples were imaged on a Talos L120C microscope at 120kV with a Ceta 16M camera. Images were analyzed in ImageJ by manually tracing around myelin and axonal membranes. The G-ratio was determined by dividing the diameter of an axon by the outer diameter of the corresponding myelin sheath. Resolution of images was sufficient to determine the major dense lines of compact myelin therefore individual lamella were then counted manually.

### Buffers and solutions

Stock Gey’s balanced salt solution (GBSS) in mM: 1.5 CaCl_2_, 0.2 KH_2_PO_4_, 0.3 MgSO_4_, 2.7 NaHCO_3_, 5 KCl, 1 MgCl_2_, 137 NaCl, 0.85 Na_2_HPO_4_, 5.6 D-glucose in sterile filtered water (0.2 µm filter pore). Culturing medium (in mM): 25% heat inactivated horse serum, 30 D-glucose, 12.5 HEPES, 25% Hank’s balanced salt solution (Thermo Fisher Scientific 24020091), in minimum essential medium with L-glutamine (Thermo Fisher Scientific, 11575032) at 320 mOsm and pH of 7.2, sterile filtered (0.2 µm). Recording artificial cerebrospinal fluid (ACSF) was composed of: (in mM) 125 NaCl, 25.0 NaHCO_3_, 1.25 mM NaH_2_PO_4_, 3.0 KCl, 25 Glucose, 2 CaCl_2_ and 1 MgCl_2_ 310 ± 5 mOsm and pH of 7.4, when perfused with 95% O_2_ and 5% CO_2_ (carbogen),

### Electrophysiological recordings

Whole-cell patch-clamp and LFP recordings were made with a MultiClamp 700B amplifier (Molecular Devices LLC., San Jose, CA, USA). Borosilicate glass pipettes (o.d. 1.5, i.d. 0.86 mm, Harvard Apparatus, Edenbridge, Kent, UK) were pulled to an open-tip resistance of 4–7 MΩ for whole-cell or <1 MΩ for LFP recordings. For whole-cell recordings, patch pipettes were filled with an internal solution containing (in mM): 135 K-gluconate; 20 KCl; 10 Na_2_-phosphocreatine; 10 HEPES; 0.1 EGTA; 2 MgATP; 0.3 Na_2_-ATP, adjusted to 302 mOsm and pH of 7.28). A liquid junction potential for this intracellular solution was corrected by –13 mV and applied to all membrane potentials reported. Some recording pipettes were supplemented with 0.5% w/v biocytin (Sigma-Aldrich, B4261) for reconstructions. The LFP recordings were made with glass pipettes filled with standard extracellular ACSF. For cell-type targeted recordings slices were illuminated with near-infrared oblique illumination, and 2P excitation of fluorescence (*Hexb*^TdTom^ or eFYP labelled *Rbp4*-Cre neurons) was digitally overlaid. Signals were analogue low-pass filtered at 10 kHz (Bessel) and digitally sampled at 50 kHz using an A-D converter (ITC-18, HEKA Elektronik Dr. Schulze GmbH, Germany) and the data acquisition software Axograph X (v.1.5.4, Axograph, RRID:SCR_014284, NSW, Australia). Bridge-balance and capacitances were fully compensated in current clamp mode. Recordings were conducted at constant 1–2 mL/min flow of carbogen-bubbled recording solution (see live imaging) at 30–33 °C.

### Statistics

All statistical comparisons were made using GraphPad Prism 10 software (GraphPad Software, Inc. Version 10.3.0., RRID SCR_002798). Datasets were tested for normality with using D’Agostino & Pearson or Shapiro-Wilk tests and comparisons between multiple groups were made using one-way ANOVA with either Tukey’s or Dunnett’s T3 post-hoc test. To test interactions between two variables, a two-way ANOVA with Tukey’s post-hoc test was used. Comparisons between OSC and *in vivo* myelin ultrastructural metrics were made using Mann-Whitney tests. A difference was considered statistically significant when *P* < 0.05. Data on graphs depicting whole slice development across weeks (**Figure 2, 4**) are displayed as ± SD and all other data on graphs are displayed as ± SEM. To ensure reproducibility all experiments were done in slice cultures were made from at least 3 animals and randomly assigned to groups with the exception EM data, which was derived from one mouse and one slice culture.

## Acknowledgements

The authors are thankful to Naomi Petersen (NIN–KNAW) and the Electron Microscopy Centre Amsterdam, Prof. Dr. med. Martin Kerschensteiner (University of München) for sharing the AAV-Mbp:mem-EGFP plasmid and Dr. Koen Kole for producing the virus. We are indebted to the research support including funding from the Institute of Chemical Immunology (project ICI0000030 and ICI000032) and grants of the Friends Foundation of the Netherlands Institute of Neuroscience (Stichting Hersenvrienden).

---

**Supplemental figure 1.**
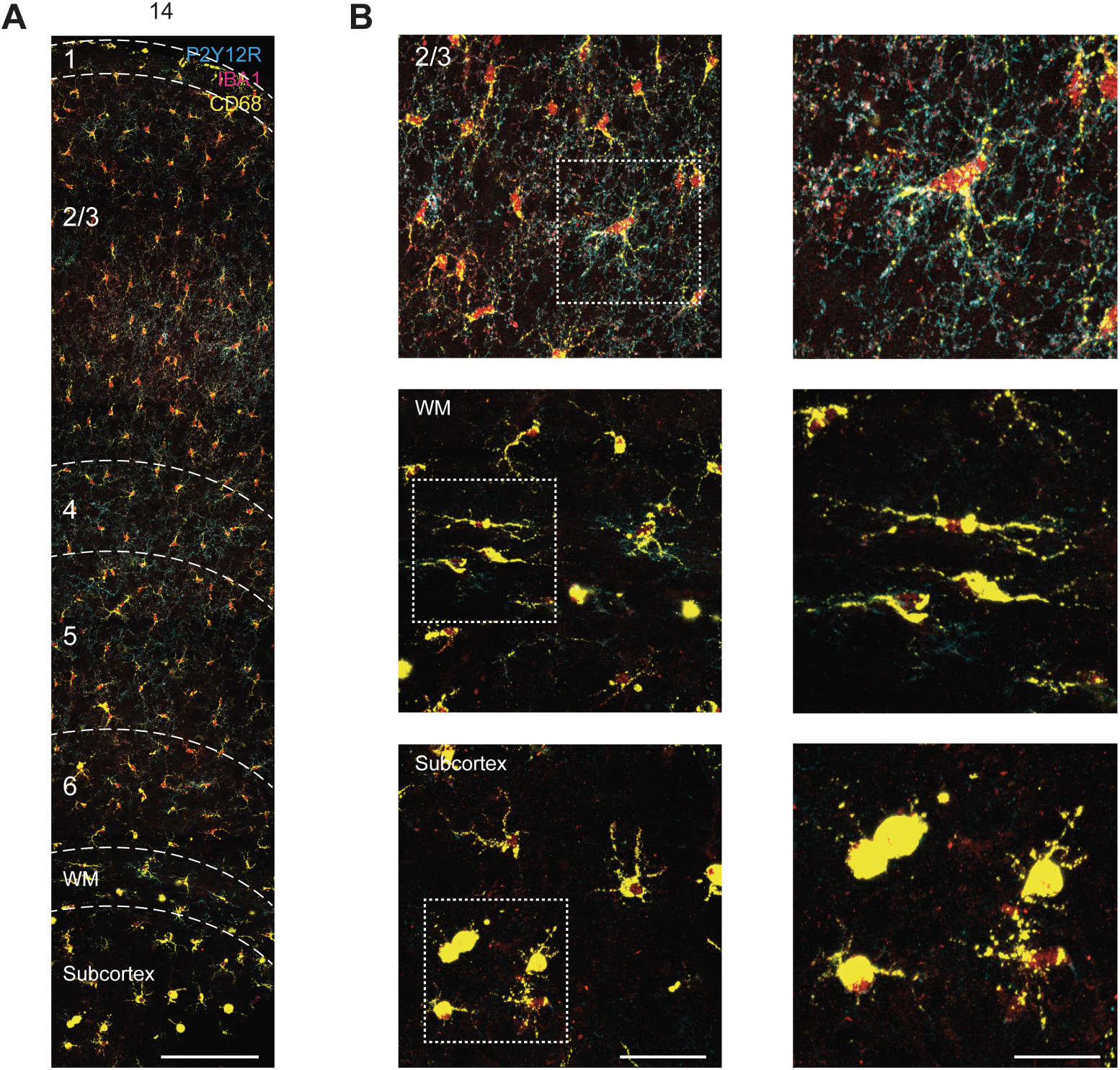
Extracortical and white-matter associated microglia are phagocytic. **A.** *Z*-projected confocal image of white matter and subcortical region from DIV 14 slice stained for P2Y12R, IBA1, CD68. Scale bar, 150 µm. **B.** (*Top*) Example image of microglia from L2/3. (*Middle*) Example image of WM associated microglia in DIV 14 slice. (*Bottom*) Example image of subcortical amyloid microglia. (Left) Scale bar 50 µm (Right) Scale bar, 25 µm.

**Supplemental Figure 2.**
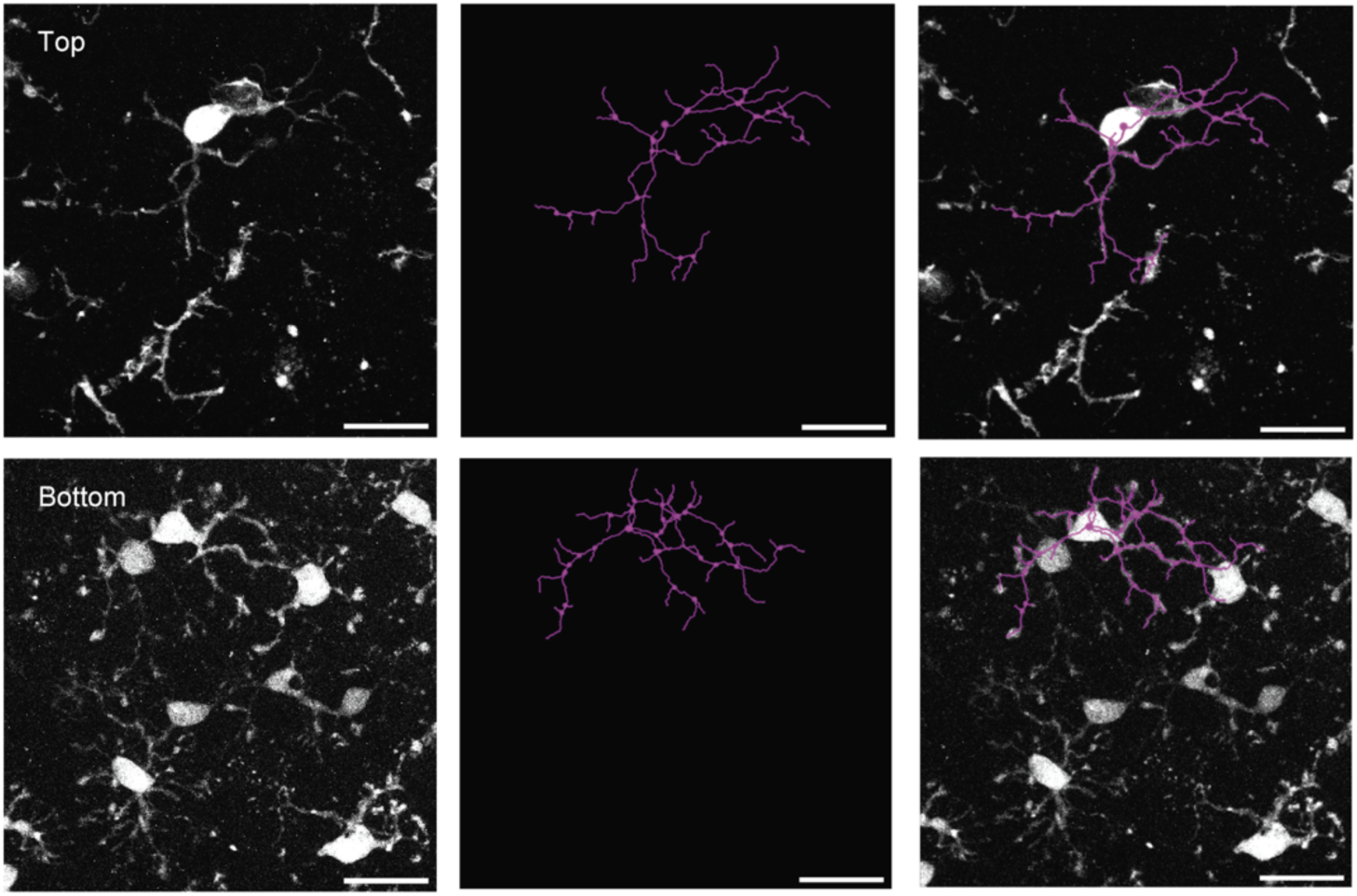
Microglia at the slice borders display a simplified morphology. Microglia from the top or bottom of the slice (top and bottom rows, respectively) displayed as 2P live images, 3D reconstructions, and merges of the two (left, center, and right column, respectively).

**Supplemental figure 3.**
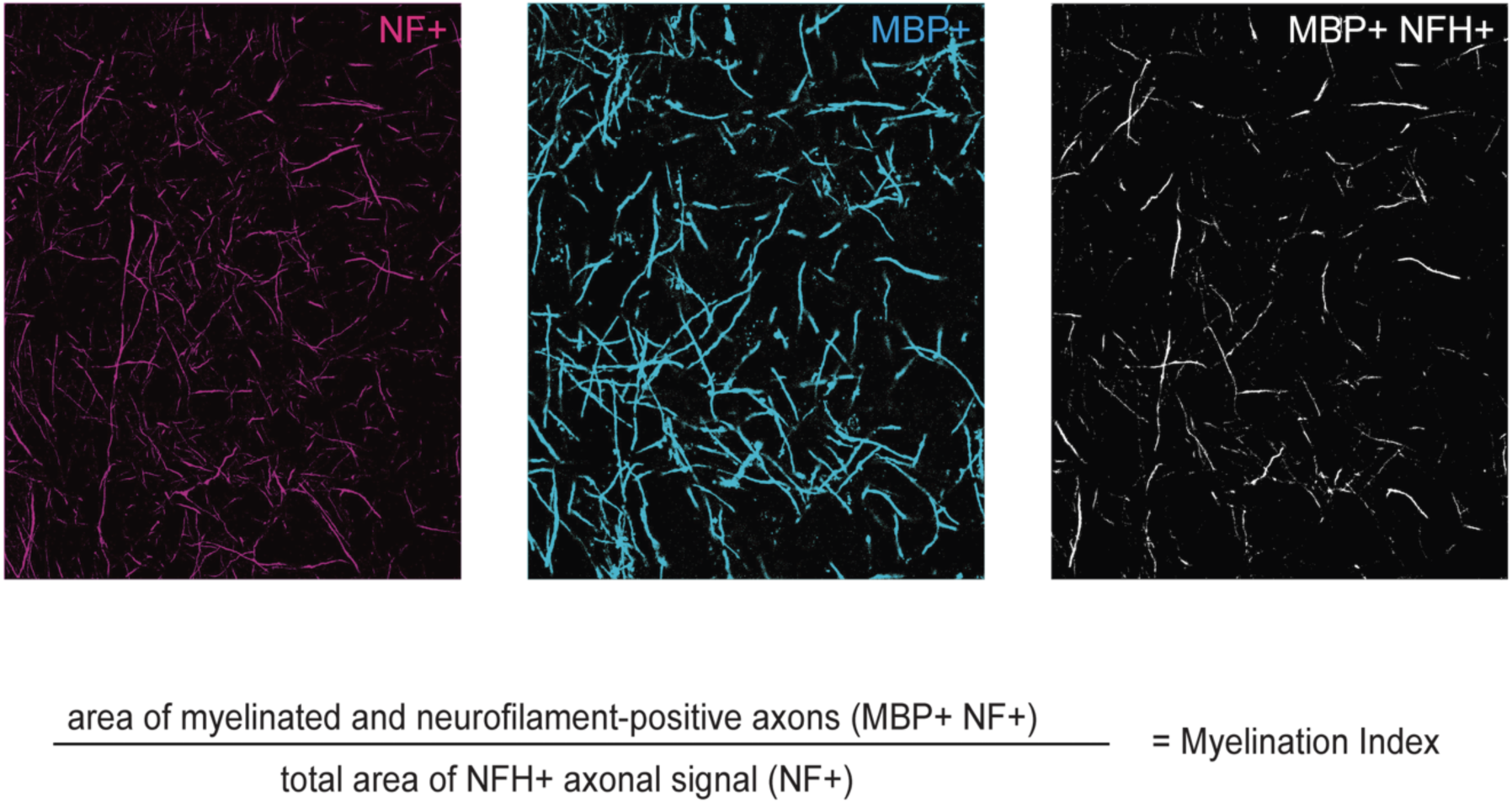
Myelination index. Example images of myelination index quantification methodology. Binarized images of area positive for NF, MBP, and MBP+NF+ were generated for DIV 7-28. Myelination index was determined by dividing area of myelinated neurofilament positive axons by total area of neurofilament positive axonal area. Scale bar 50 µm

**Supplemental figure 4.**
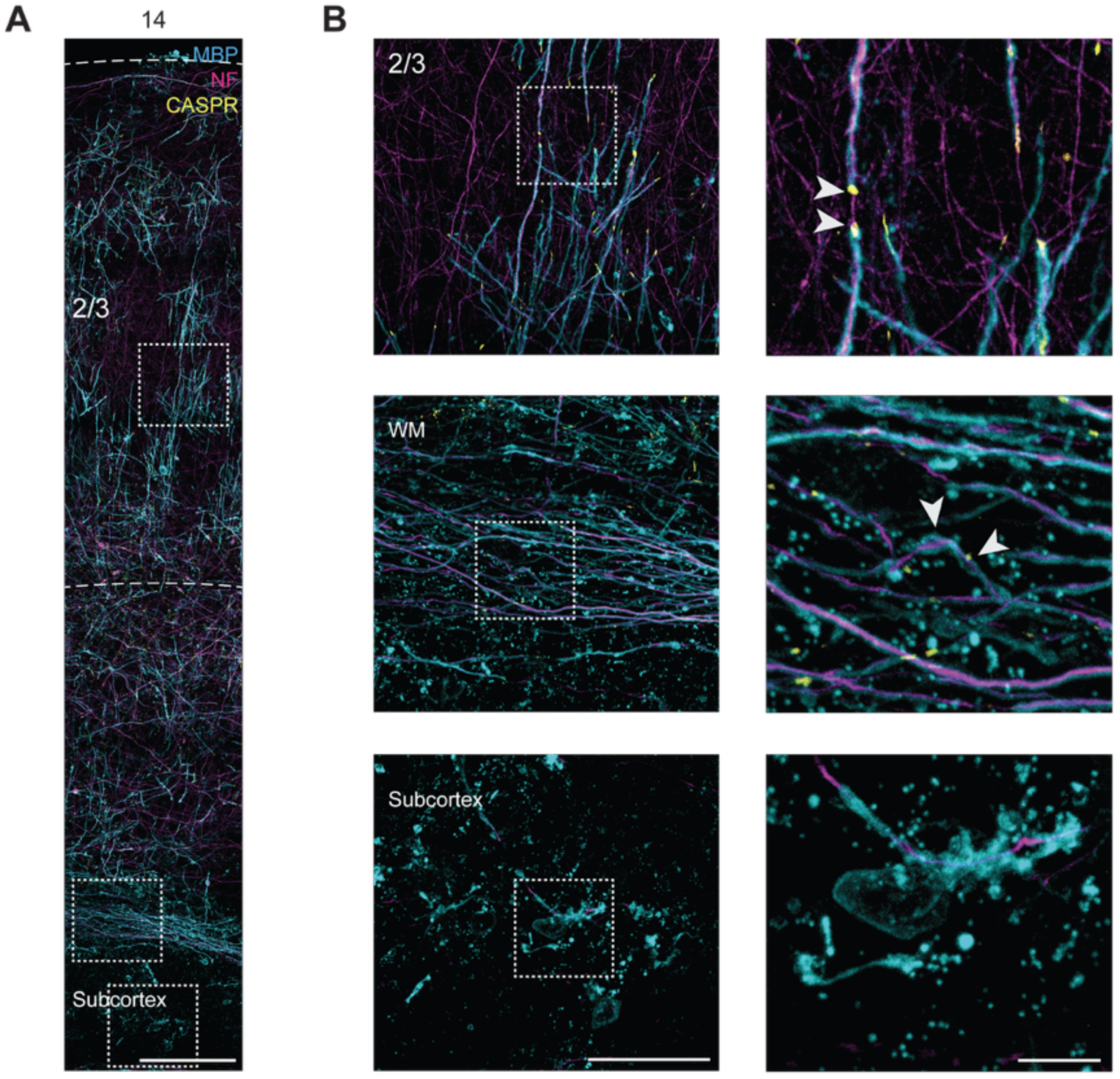
White matter and subcortical myelin degeneration. **A.** *Z*-projected confocal image of white matter and subcortical region from DIV 14 slice stained for MBP, NF, and CASPR. Scale bar 150 µm. **B.** (*Top*) Example image of patchy myelinated axons from L2/3. (*Middle*) Example image of WM degeneration in DIV 14 slice. White arrows indicate myelin swellings. (Bottom) Example image of subcortical degeneration with myelin debris. (*Left*) Scale bar 50 µm (*Right*) Scale bar 10 µm

**Supplemental figure 5.**
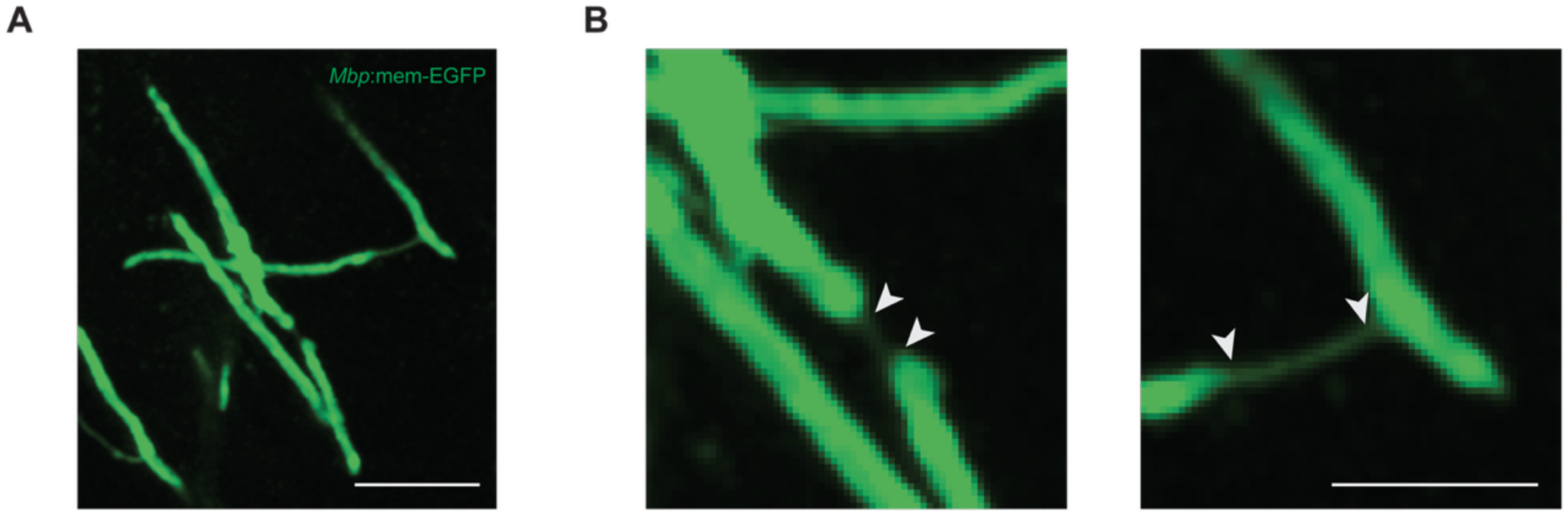
Paranodal bridge. **A.** Z-projected live two photon image of AAV2/1-*Mbp*:mem-EGFP labelled myelin sheaths linked by paranodal bridges. Scale bar 20 µm. **B.** Example images of two forms of paranodal bridges indicated by white arrows. (Left) A paranodal bridge linking two myelin sheaths along the same putative axon. (Right) T-shaped paranodal bridge linking putatively disparate axons. Scale bar 10 µm.

**Supplemental Table 1.**

| Primary antibody | Supplier, cat. # | Concentration | RRID |
| --- | --- | --- | --- |
| Neurofilament-H phospho (NFH) - mouse monoclonal | BioLegend, 801601 | 1:200 | AB_2564641 |
| Caspr (Contactin-associated protein) - rabbit polyclonal | Abcam, ab34151 | 1:500 | AB_869934 |
| Myelin basic protein (MBP) - rat monoclonal | Millipore (Merck), MAB386 | 1:200 | AB_94975 |
| Green fluorescent protein (GFP) - chicken polyclonal | Abcam, ab13970 | 1:500 | AB_300798 |
| Ionized calcium-binding adaptor molecule 1 (Iba1) - guinea pig recombinant monoclonal | Synaptic Systems, 234 308 | 1:500 | AB_2924932 |
| COUP-TF-interacting protein 2 (CTIP-2) – rat monoclonal | Abcam, ab18465 | 1:400 | AB_2064130 |
| Neurofilament H non-phosphorylated (SMI-32) – mouse monoclonal | BioLegend (Covance), SMI-32P | 1:1000 | AB_10719742 |
| Purinergic receptor P2Y12 – rabbit polyclonal | Anaspec, AS-55043A | 1:250 | AB_2298886 |
| Cluster of differentiation 68 (CD68) | Abcam, ab53444 | 1:250 | AB_869007 |

| Secondary antibody | Supplier, cat. # | Concentration | RRID |
| --- | --- | --- | --- |
| Goat anti-mouse IgG/IgM (H+L) F(ab')<sub>2</sub>, Alexa Fluor 488 | Invitrogen/Thermo Fisher, A-10684 | 1:500 | AB_2534064 |
| Goat anti-rabbit IgG (H+L) highly cross-adsorbed, Alexa Fluor 568 | Invitrogen/Thermo Fisher, A-11036 | 1:500 | AB_10563566 |
| Goat anti-rat IgG (H+L) cross-adsorbed, Alexa Fluor 633 | Invitrogen/Thermo Fisher, A-21094 | 1:500 | AB_2535749 |
| Goat anti-chicken IgY (H+L), Alexa Fluor 488 | Invitrogen/Thermo Fisher, A-11039 | 1:500 | AB_2534096 |
| Goat anti-guinea pig IgG (H+L) highly cross-adsorbed, Alexa Fluor 594 | Invitrogen/Thermo Fisher, A-11076 | 1:500 | AB_2534120 |
| Goat anti-rat IgG (H+L) cross-adsorbed, Alexa Fluor 647 | Invitrogen/Thermo Fisher, A-21247 | 1:500 | AB_141778 |
| Goat anti-mouse IgG (H+L) highly cross-adsorbed, Alexa Fluor 568 | Invitrogen/Thermo Fisher, A-11031 | 1:500 | AB_144696 |

## Notes

### Competing Interest Statement

The authors have declared no competing interest.

